# Soil metatranscriptomics: An improved RNA extraction method toward functional analysis using nanopore direct RNA sequencing

**DOI:** 10.1101/2022.11.20.517272

**Authors:** Abdonaser Poursalavati, Vahid J. Javaran, Isabelle Laforest-Lapointe, Mamadou L. Fall

## Abstract

Soil microbes play an undeniable role in sustainable agriculture, plant health, and soil management. A deeper understanding of soil microbial composition and function has been gained through next-generation sequencing. While soil metagenomics has provided valuable information about microbial diversity, issues stemming from RNA extraction, low RNA abundance in some microbial populations (e.g., viruses), and mRNA enrichment have slowed the progress of soil metatranscriptomics. A variety of soil RNA extraction methods have been developed so far. Yet none of the available protocols can obtain RNA with high quality, purity, and yield for third-generation sequencing. This latter requires RNA with high quality and large quantities (with no or low contamination, such as humic acids). Also, use of commercial kits for in-batch soil RNA extraction is quite expensive, and these commercial kits lack buffer composition details, which prevents the optimization of protocols for different soil types. An improved and cost-effective method for extracting RNAs from mineral and organic soils is presented in this paper. An acidic sodium acetate buffer and phosphate buffer with modifications to bead-beating and nucleic acid precipitation lead to higher RNA yields and quality. Using this method, we obtained almost DNA-free RNA. By using nanopore’s direct RNA sequencing, the extracted contamination-free RNAs were successfully sequenced. Lastly, taxonomic groups such as bacteria, fungi, archaea, and viruses were classified and profiled as well as functional annotation of the datasets was carried out using an in-house customized bioinformatics workflow.

## Introduction

As a highly dynamic ecosystem, the soil is a reservoir of various microbes such as bacteria, archaea, viruses, fungi, and protozoa (Jansson and Hofmockel 2020; Sharuddin et al. 2022). To balance the negative effects of intensive agriculture on soil functions, we rely on the synergistic abilities of microorganisms to regulate the biogeochemical cycle in the environment (Cavicchioli et al. 2019; Sharuddin et al. 2022). Yet, we lack a deep understanding of how soil microbial community composition and functions support and influence soil biogeochemical processes. A variety of environmental DNA (eDNA) sequencing methods have been employed to gain unprecedented insight into soil microbial community composition and diversity (Podolyan and Grelet 2021; Wang et al. 2009c). For example, amplicon-based gene sequencing (Aguiar-Pulido et al. 2016; Zuñiga et al. 2017) and shotgun metagenomics (Azeem et al. 2021; Shakya et al. 2019) are widely used for deeper investigation of the soil microbiome. However, these DNA-based methods are incapable of accurately assessing microbiome functionality and can hardly discern between active and inactive microbiome members (Sharuddin et al. 2022). Environmental RNA (eRNA) sequencing is essential to reveal the functions from active microbiome members.

Several metatranscriptomic studies have been carried out over the last decade to analyze gene expression, regulation, and pathways in many different types of biotope (Hayden et al. 2018; Rajarapu et al. 2015; Sharuddin et al. 2022). However, in the field of soil and environment metatranscriptomics, the lack of a universal approach to RNA has hampered progress. Obtaining high-quality and high-quantity RNA from environmental samples has always been a challenge, especially in soil samples. Following RNA extraction, various compounds like humic and fulvic acids, as well as polysaccharide compounds, are co-extracted with RNA (Wang et al. 2012a) and incorporated into downstream enzymatic reactions (such as restriction enzymes, probe hybridization, RNA or DNA digestion, polymerase chain reaction [PCR], reverse transcription, sequencing, and quantification) (Alm Elizabeth et al. 2000; Chaparro-Encinas et al. 2020; Wang and Fujii 2011; Zipper et al. 2003). Clay minerals, such as Ca^2+^, Mg^2+^, Fe^3+^ and Al^3+^, can also absorb a significant amount of nucleic acid molecules, resulting in a lower level of RNA extraction (Goring and Bartholomew 1952). In addition, soil samples can contain a wide range of contaminants including proteins, phenolic compounds, salts, and metal ions (Griffiths et al. 2000; Wilson 1997). In addition to the aforementioned problems, less than 5% of total RNA is mRNA in environmental samples, which is highly vulnerable to RNase degradation and has a short half-life span (Deutscher 2006; Ranjan et al. 2021; Sharma et al. 2012; Steglich et al. 2010). Therefore, the development of efficient high-quality RNA extraction protocols for different soil sample types is required. To extract RNA from a variety of soil types, different manual extraction methods have been optimized (Lever et al. 2015; Lim et al. 2016; Mettel et al. 2010; Paulin et al. 2013; Pei et al. 2021; Peršoh et al. 2008; Qin et al. 2016; Sharma et al. 2012; Thorn et al. 2019; Wang et al. 2012b). However, soil samples exhibit many heterogeneities that have hampered the development of a universal RNA extraction method. The manual extraction process has been standardized and continuously optimized over the past three decades, based on soil sample types. Although there are several commercial RNA extraction kits that have been used for soil samples, it is not possible to optimize each kit individually based on the components, since they are not disclosed. Also, under certain conditions, it may be necessary to increase the sample mass for RNA extraction from low-biomass samples to isolate low-copy-number mRNA; therefore, using commercial kits is not cost-effective and feasible (Lever et al. 2015; Lim et al. 2016; Thorn et al. 2019).

In this study, our main goal was to develop a high-yield and high-quality RNA extraction method for two types of agricultural soils (mineral and organic) for taxonomic and functional analysis of soil microbial communities. We used Griffiths’ extraction method (Griffiths et al. 2000) to form the backbone of our improved and developed extraction procedure. The most important modifications we made were the use of phosphate buffer and acidic sodium acetate buffers during the bead beating and phenol-chloroform extraction processes, respectively. It was surprising to find that the sodium acetate buffer not only decreased the humic substances, but also allowed us to extract DNA-free total RNA. We compared our developed extraction method with four more popular manual extraction methods developed for soil nucleic acids extraction. Our developed method improved not only the purity, but also the integrity of extracted RNA, which makes the extracted materials suitable for diverse kinds of molecular biology investigations at a reasonable price, ten times cheaper than using disparate commercial kits for DNA and RNA isolation. The extracted RNA was then successfully used for direct RNA nanopore sequencing. Four soil microbiome libraries were prepared from mineral and organic soils, and an overarching review of the soil microbiome was characterized after sequencing and data analysis.

## Materials and methods

### Soil sampling and analysis of soil physicochemical properties

Four mineral soil horizons of 5 to 15 cm depth were sampled from Agriculture and Agri-Food Canada’s experimental farm (vineyard) in Frelighsburg, Quebec, Canada. As well, four organic soil horizons were sampled at a depth of 0 to 10 cm from Agriculture and Agri-Food Canada’s experimental farm (lettuce field) in Sainte-Clotilde, Quebec, Canada. The soil samples were stored at −20 °C until they were used. Our soil samples were submitted to AgroEnviroLab for the determination of soil pH and for the extraction and quantification of main exchangeable minerals, aluminum (Al), copper (Cu), zinc (Zn), manganese (Mn) and iron (Fe), through the Mehlich 3 extraction method (Mehlich 1984).

### Total RNA extraction by improved method

A decontaminating mixture of sodium hypochlorite, 10% v/v; sodium dodecyl sulfate (SDS), 1% w/v; sodium hydroxide (NaOH), 1% w/v; and sodium bicarbonate (NaHCO3), 1% w/v, was made to clean up the work surfaces (Fischer et al. 2016). To eliminate nucleases, all implements and tools were autoclaved for 15 minutes at 121 °C. In addition, the water stock used for RNA elution was treated by RNASecure (Ambion) to reduce the possibility of nuclease contamination during extraction.

The samples were passed through a 4-mm mesh sieve, and RNA was extracted directly from them (Fig. 1). For mineral and organic soil, 250- and 200-mg stone-free soil samples were collected and transferred into 2-ml screw-tubes, respectively. Each tube contained 1 gram of 0.1-millimeter silica beads and three beads of 0.3 millimeter size. Next, 200 μl pre-heated (60 °C) extraction buffer (10% Cetyltrimethyl ammonium bromide [CTAB], 0.7 M NaCl, 3.4% Polyvinylpyrrolidone [PVP], 240 mM phosphate buffer, pH = 5.8), 400 μl 150 mM phosphate buffer (pH = 5.8), 10 μl 2-Mercaptoethanol, 300 μl water-saturated phenol, and 200 μl Chloroform were added. Before the bead-beating steps, tubes were incubated on ice and cooled for 2 minutes. Two FastPrep bead-beating steps (6.5 m/s for 20 seconds) and one intermediate cooling step on ice (for 1 min) were completed. After that, samples were vortexed for 5 min periodically, and during vortexing time, tubes were placed on ice to keep them cooled. After the centrifugation at 10,000X g for 2 min at 4 °C, the aqueous phase (approximately 550 ul) was transferred into a new 2-ml tube. Next, 350 μl 3M NaAc (pH = 4.6) was added to the previous solution and incubated on ice for 2 min. Then, 600 μl water-saturated phenol was added, mixed, and incubated on ice for 3 min. Next, 300 μl chloroform was added and mixed vigorously. After incubation on ice for 3 min, tubes were centrifuged for 10 min at 10,000X g at 4 °C. The aqueous phase was transferred into a new 2-ml tube, and another chloroform extraction (1 v/v) was performed and centrifuged for 10 min at 10,000X g at 4 °C.

**Fig. 1.**
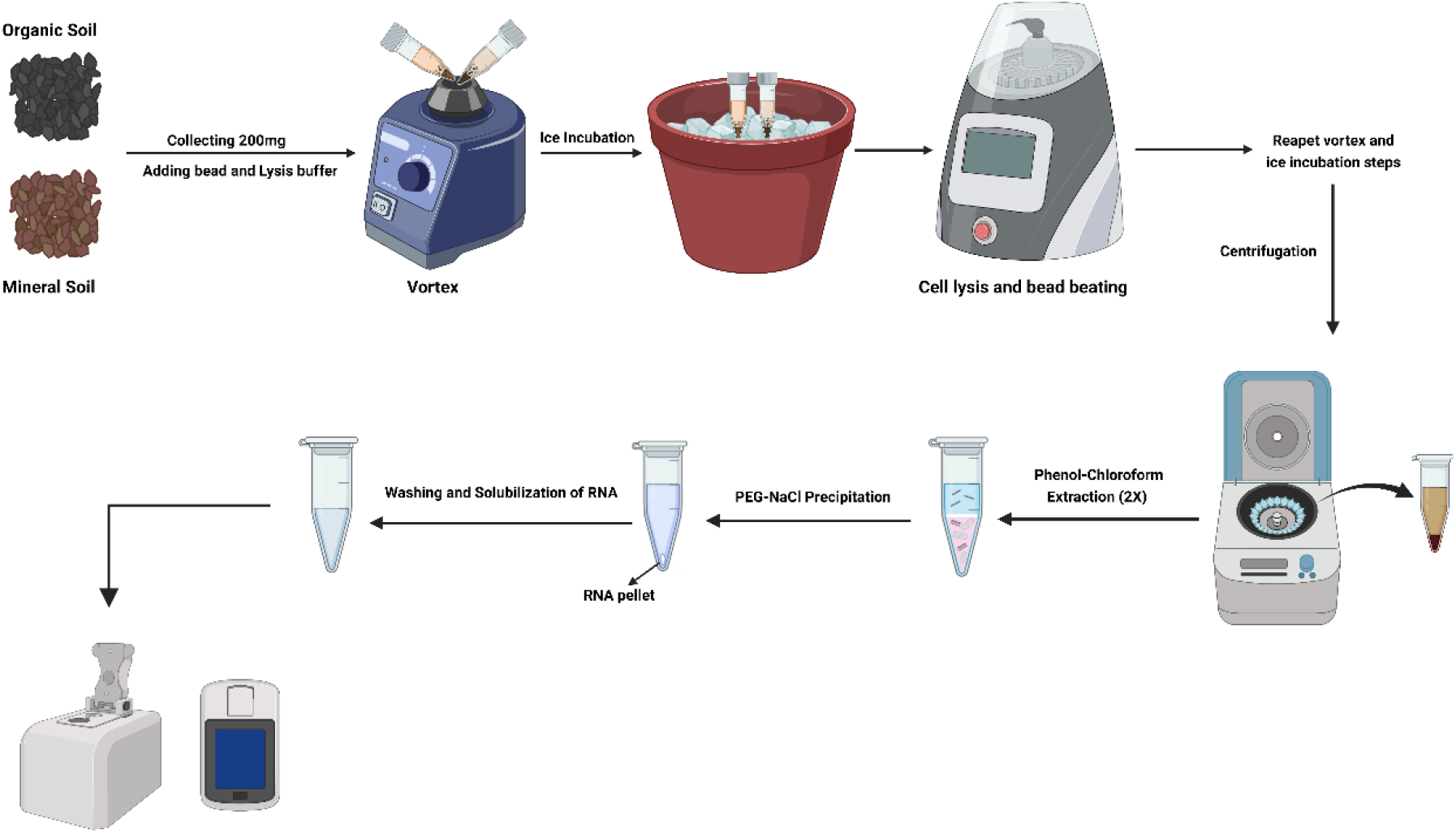
The workflow of total RNA extraction from two types of soils (mineral and organic) using the improved RNA extraction method

**Fig. 2.**
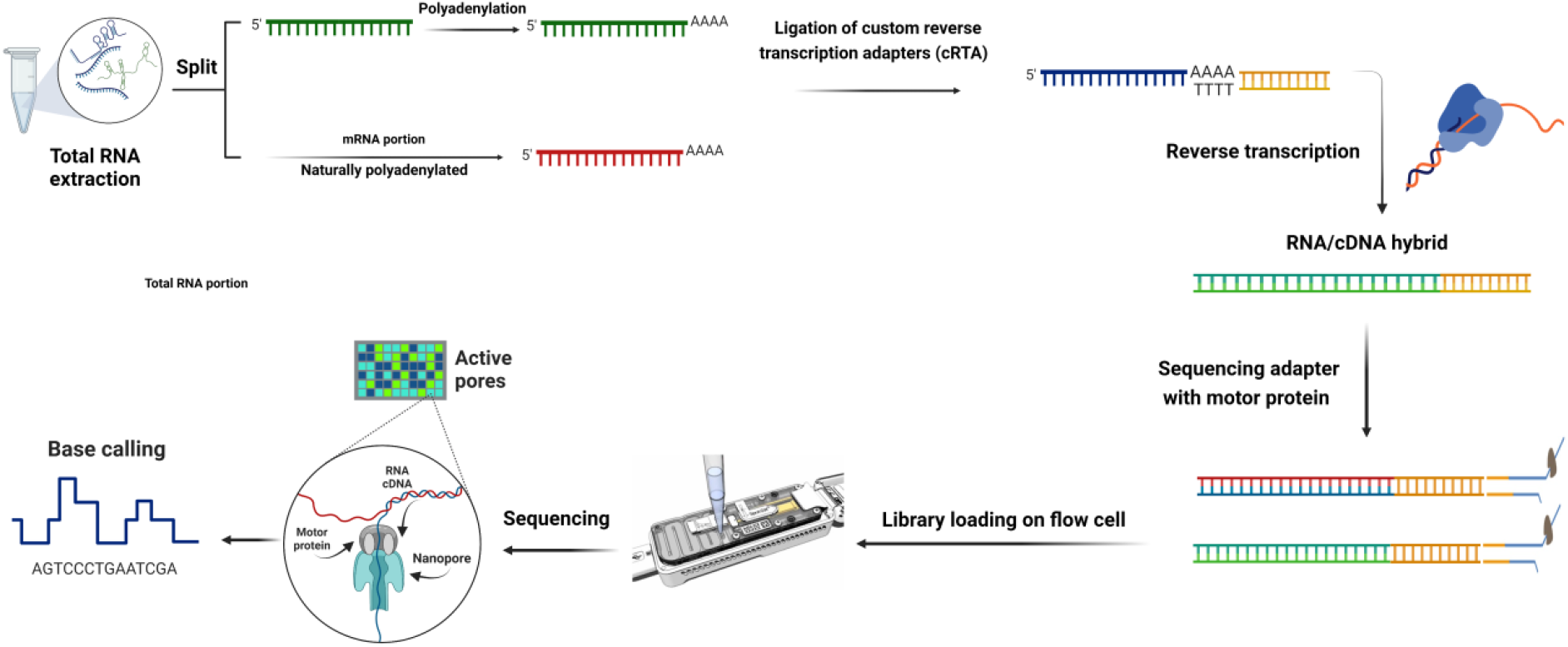
The library and sequencing workflow of direct RNA nanopore sequencing for organic and mineral soil samples. Ligation is done in two steps: first, cRTA adapters are ligated to all poly-A sequences, then cDNA strand is synthesized to form hybrid RNA/cDNA. In the second stage, the sequencing adapter (along with the motor protein) is ligated to the hybrid sequences. When the motor protein is connected to the nanopore protein, only the RNA strand passes through the pore, and the native RNA is sequenced.

In the next step, the aqueous phase (approximately 500 to 600 ul) was transferred into a new 1.5-ml tube. One volume PEG-NaCl precipitation buffer (0.6 M NaCl and 30% PEG-8000) was added, and the solution was incubated on ice in a refrigerator (4 °C) for 20 min. To visually monitor the RNA pellet precipitation, 1 μl glycogen RNA grade can be added. The tubes were centrifuged for 20 min at 15,000 rpm at 4 °C. After that, the PEG supernatant was removed carefully by pipetting. The RNA pellet was washed with 80% cold ethanol two times. After each washing step, a centrifugation step at 15,000 rpm for 5 min at 4 °C was done. Finally, the RNA pellet was dried under laminar hood and dissolved in 50 μl treated water with RNAsecure.

### Total RNA extraction by other conventional methods

In this study, Griffiths’ (Griffiths et al. 2000), Thorn’s (Thorn et al. 2019), Sharma’s (Sharma et al. 2012), and Angel’s (Angel et al. 2012) RNA extraction methods, which are the main methods for RNA extraction from soil and constitute the backbone of our improved and optimized method, were applied in the same eight soil samples described in the previous section to compare their efficiencies with our improved method.

### Quality and quantity evaluation of extracted RNA

RNA extracted using the five different methods was electrophoresed on 1.0% agarose gel to confirm its integrity and quality. Nanodrop 2000 spectrophotometry (Thermo Fisher Scientific, Waltham, MA) was used to determine the purity of the extracted RNA. Pure RNA was defined as having the A260/A280 ratio = 1.8 ~ 2.2 and the A260/A230 ratio > 1.8. Various wavelengths (320 or 270 nm) and spectrum ratios (from 465 to 665) have been used to quantify humic substances (Wnuk et al. 2020). Mettel *et al*. concluded that 400-nm wavelengths were more useful than 320-nm measurements for quantifying humic substances due to the avoidance of the overlapping absorbance errors that occurred in the absorbance spectrum of RNA and humic acids at 320-nm measurements (Mettel et al. 2010). Hence, 400-nm wavelength was considered for humic acid measurements in this study. The quantification of DNA and RNA at different steps of extraction and library preparation were performed by qubit spectrofluorometry using Qubit dsDNA High Sensitivity and Qubit™ RNA High Sensitivity kits, respectively (Qubit Fluorometer, Invitrogen, LifeTechnologies).

### DNase treatment and polyadenylation

According to gel electrophoresis results, no DNA contamination was observed in extracted RNAs. However, to be sure, total extracted RNAs were treated with TURBO DNase (Ambion) according to the manufacturer’s instructions. In the presence of 1X DNase buffer and 10 units of TURBO enzyme, the reaction solution was incubated for 30 mins at 37 °C, then phenol-chloroform extraction was used to remove the DNase enzyme from the treated sample.

To prepare and sequence libraries, two groups of extracted RNA for each soil type were defined: mRNA group (naturally polyadenylated messenger RNA) and total RNA group (mRNA + non-polyadenylated RNA). Because reverse transcription of the direct RNA nanopore sequencing kit is based on ligation of pre-annealed double-stranded adapters (containing poly-T sequence), the total RNA group was polyadenylated as followed, whereas the mRNA group was not polyadenylated. Next, 10 ug total RNA, 1 μl (five units) *E. coli* Poly(A) Polymerase (NEB# M0276), 2 μl 10X *E. coli* Poly(A) Polymerase Reaction Buffer, 2 μl Adenosine triphosphate (ATP), and 1 μl (40 unit) RNasin^®^ Plus Ribonuclease Inhibitor (Promega Corporation) in a 20-μl reaction solution were incubated at 37 °C for 3 mins. After that, polyadenylated RNAs were purified from enzyme reactions using the RNAClean XP bead purification system according to the manufacturer’s instruction.

### Direct RNA nanopore sequencing

The SQK-RNA002 kit (Oxford Nanopore Technologies) was used to generate direct RNA sequencing libraries according to the manufacturer’s protocol with some modifications (2). Since this kit does not include commercial barcodes for multiplexing samples and sequencing them simultaneously on a single flow cell, four custom reverse transcription adapters (cRTA) based on the DeePlexiCon protocol (Smith et al. 2020) were used for multiplexing.

In summary, 500 ng of polyadenylated RNA was ligated with cRTA using T4 DNA ligase (New England Biolabs) in the presence of NEBNext Quick Ligation buffer (New England Biolabs) for 15 minutes at room temperature. The single-stranded cDNA was synthesized with SuperScript III (Thermo Fisher Scientific) for 50 minutes at 50 °C, followed by the inactivation of the enzyme reaction for 10 minutes at 70 °C. RNA/cDNA hybrids were then purified using Agencourt RNAClean XP beads. We measured the quantity of each library with the Qubit dsDNA High Sensitivity kit and then pooled the libraries by considering an equal amount of each. The RMX adapter (sequencing adapter) was ligated into the RNA/cDNA hybrid complex using T4 DNA ligase for 15 minutes at room temperature, and then the pooled library was purified with Agencourt RNAClean XP beads. Final quantification of the pooled library was performed using the Qubit dsDNA High Sensitivity kit, followed by loading on the R9.4.1 flow cell connected to the MinION Mk1B Sequencing Instrument (Oxford Nanopore Technologies).

### Bioinformatics analysis pipeline

In high-accuracy mode, base-calling and quality assessment of sequencing data were performed using Guppy (Oxford Nanopore, v6.1.7). We discarded reads with a poor quality score (<7) and lengths under 100 nucleotides. For demultiplexing, DeePlexiCon was used with default settings. Then, to trim barcodes and middle adapters, Guppy barcoder default parameters were used. Next, NanoPlot (v1.40.0) (De Coster et al. 2018) was used to generate direct RNA sequencing metrics such as N50, read count, and quality score (Q). Seqkit (Shen et al. 2016) fq2fa option was used to convert the trimmed fastq files to fasta files. RATTLE pipeline with a reference-free algorithm (de la Rubia et al. 2022) was used for read clustering, error correction of reads, and read polishing. From each cluster, a consensus transcript was extracted, and the abundance was determined by counting the total reads in each cluster. With SortMeRNA (Kopylova et al. 2012), ribosomal RNA was then extracted from total RNA samples (from barcodes 1 and 2) using its sensitive database (smr_v4.3_sensitive_db). Kraken2 v2.0.8-beta (Wood et al. 2019) was used to classify 16S and 18S rRNA reads using the SILVA database (Quast et al. 2013). The non-ribosomal reads (from barcodes 1 and 2), as well as the mRNA reads (from barcodes 3 and 4), were taxonomically classified by Kraken2 using the PlusPF database. Pavian v1.0 (Breitwieser and Salzberg 2020) and Recentrifuge (Martí 2019) were used for comparative analysis of taxonomical classification results. After Kraken2 analysis, clusters without hits were extracted, and these clusters were analyzed taxonomically with Centrifuge 1.04 (Kim et al. 2016) using a custom-built database, whose sequences were obtained from NCBI RefSeq (Virus, Bacteria, Archaea, SAR, Protozoa, and Fungi, access date 03.06.2022) using Genome_updater 0.5.1 (https://github.com/pirovc/genome_updater). The minimum hit length (MHL) of 50 was applied on centrifuge hits (Kim et al. 2016).

Through Trinotate v3.2.2 (https://github.com/Trinotate/Trinotate.github.io), BLASTX and BLASTP searches (Altschul et al. 1990) were conducted on the SwissProt database (Bairoch and Apweiler 1997) to analyze the functional annotation of transcripts. EggNOG-mapper version 2.1.6 (Cantalapiedra et al. 2021; Huerta-Cepas et al. 2019) was used to obtain EggNog, KEGG (Kanehisa and Goto 2000), and COG annotations (Tatusov et al. 2000). SigmaPlot v14.5 (Systat Software, San Jose, CA) was used to visualize the results of functional annotation. A GO Slim metagenomics database was then substituted for the pre-packed GO Slim database in the Trinotate package (Poursalavati et al. 2021). Using Trinotate_GO_to_SLIM, gene ontology identifiers were assigned to clusters. The annotated KEGG orthologs (KOs) were then processed and analyzed using the FuncTree 2, an automated annotation server (Darzi et al. 2019). A summary of the bioinformatics analysis steps is shown in Fig. 3.

**Fig. 3.**
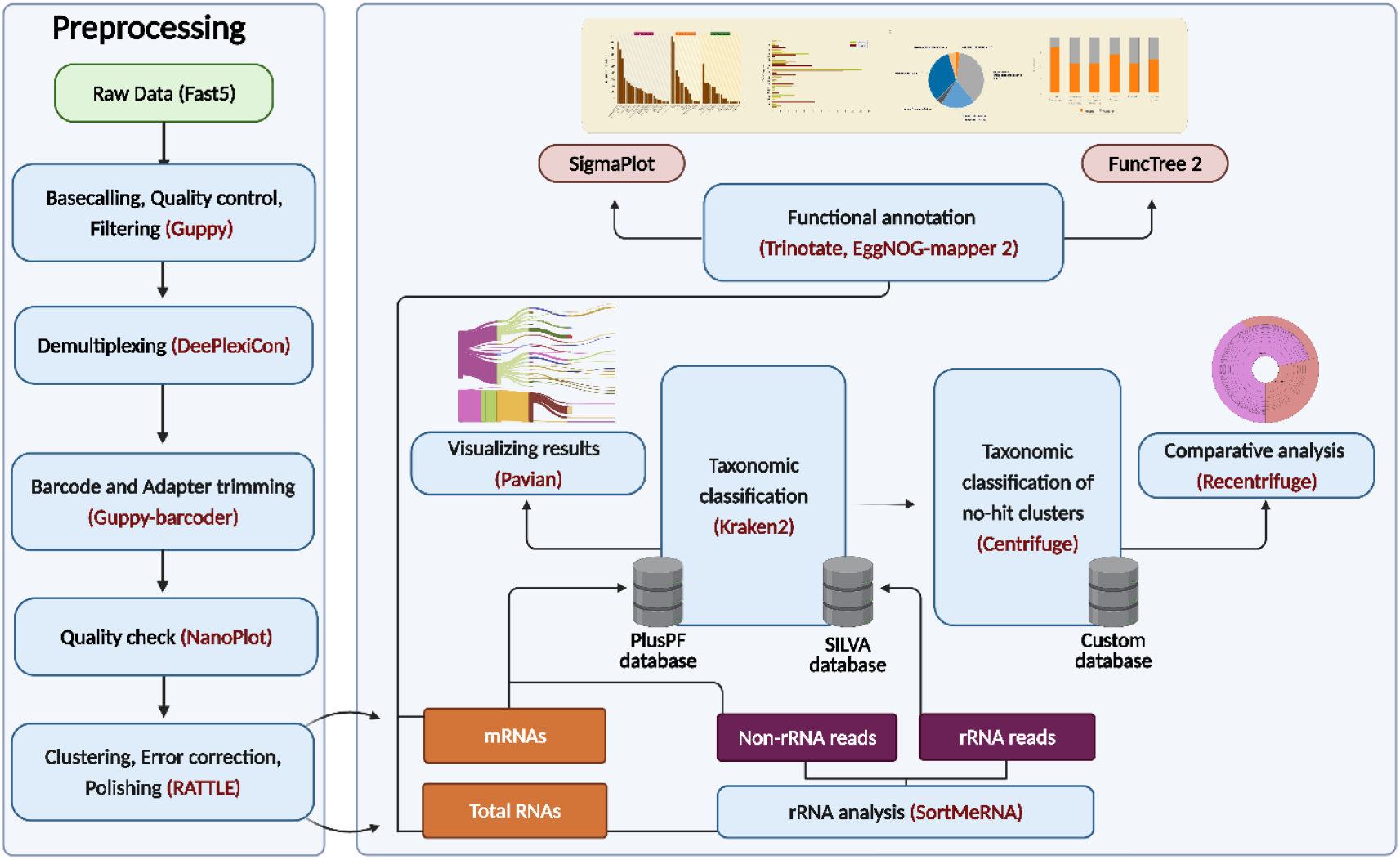
Metatranscriptomic data analysis workflow for direct RNA nanopore sequencing data. Following preprocessing and demultiplexing of the data, RATTLE pipeline was used for read clustering, error correction, and creating consensus transcripts for each cluster. Using SortMeRNA, ribosomal RNA was identified, and then rRNA-free data and ribosomal reads were separately analyzed with Kraken2 (via PlusPF and SILVA databases). Clusters without hit were extracted and further analyzed using custom-built databases with the Centrifuge tool. Recentrifuge and Pavian were used to visual inspection of taxonomy analysis. A functional annotation of the data was performed using Trinotate and EggNOG-mapper 2, and the data was visualized using FuncTree 2 and SigmaPlot.

## Results

### Soil physiochemical properties

The adsorption of RNA to clay minerals is one of the main problems in soil RNA extraction (Hashizume 2015). Prior to RNA extraction, we analyzed the physicochemical properties of collected samples of mineral and organic soils. Texture, organic content, and cations were widely different in mineral and organic soil samples. Mineral soil samples were found to have acidic pH values in the range of 5.2 to 5.6, while organic soils were in the range of 6 to 6.8. In terms of organic content, mineral soil samples ranged from 5.2% to 5.5% and, on the other hand, organic soils were 67.6% to 73.7% (**Table 1**).

**Table 1.** Physicochemical characteristics of mineral and organic soil samples

| Properties | Units | Mineral soils |  |  |  | Organic soils |  |  |  |
| --- | --- | --- | --- | --- | --- | --- | --- | --- | --- |
|  |  | S1 | S2 | S3 | S4 | S1 | S2 | S3 | S4 |
| pH | - | 5.6 | 5.5 | 5.4 | 5.2 | 6.5 | 6.0 | 6.4 | 6.8 |
| Organic content | % | 5.4 | 5.5 | 5.4 | 5.2 | 72.3 | 73.7 | 71.7 | 67.6 |
| Organic carbon | % | 4.8 | 4.8 | 4.8 | 4.8 | 45.2 | 45.2 | 45.2 | 45.2 |
| CEC | meq/100g | 16.7 | 17.0 | 17.6 | 15.4 | 31.0 | 36.4 | 34.4 | 34.6 |
| P | kg/ha | 72 | 56 | 39 | 39 | 38 | 49 | 35 | 26 |
| K | kg/ha | 101 | 98 | 133 | 156 | 208 | 279 | 236 | 129 |
| Ca | kg/ha | 3236 | 3046 | 2697 | 1839 | 10636 | 10532 | 11069 | 12356 |
| Mg | kg/ha | 142 | 135 | 169 | 106 | 668 | 719 | 670 | 646 |
| Al | ppm | 994 | 1042 | 1143 | 1176 | 26 | 29 | 41 | 51 |
| Mn | ppm | 49.2 | 39.7 | 39.0 | 36.4 | 5.4 | 5.4 | 5.0 | 5.5 |
| Cu | ppm | 2.29 | 2.02 | 2.41 | 3.45 | 3.25 | 5.26 | 3.94 | 3.34 |
| Zn | ppm | 2.27 | 2.08 | 2.13 | 2.04 | 9.41 | 15.88 | 41.42 | 77.62 |
| B | ppm | 0.43 | 0.33 | 0.39 | 0.38 | 1.20 | 1.24 | 1.34 | 1.47 |
| Fe | ppm | 193 | 194 | 164 | 188 | 199 | 196 | 178 | 164 |

### Optimization of RNA extraction method for mineral and organic soil samples

The Griffiths’ (Griffiths et al. 2000), Thorn’s (Thorn et al. 2019), Sharma’s (Sharma et al. 2012), and Angel’s (Angel et al. 2012) CTAB extraction and PEG precipitation methods were the backbone of our optimized method for RNA extraction. Cells were lysed in the first step by choosing different-sized beads and changing the bead beating speed and duration, but the main change was adding a phosphate buffer into the sample based on measured pH value (Guerra et al. 2020). A pre-experiment, which used a variety of pH ranges (4.3 to 8) was designed to decrease the amount of co-extracted humic substances by considering the lowest adsorption of RNA to the soil. We found that the amount of humic substances was reduced optimally at pH 5.8 (Supplementary Fig. 1). It was observed that the amount of RNA extracted was the same at pH 5.5, 5.8, and 6, but the lowest levels of humic substances were found at pH 5.8. While lower pH ranges (<5) are recommended in the literature for removing humic substances (Mettel et al. 2010), we noticed that most of the nucleic acids in our experiment were absorbed at pH < 5.5. A second phenol-chloroform extraction in an acidic condition was considered in this protocol to isolate DNA-free RNA and reduce remaining humic compounds. A low-pH (4.6) sodium acetate buffer was used to create the acidic condition within the aqueous phase from the previous step. Sodium acetate has been used previously to separate DNA from RNA (Chomczynski and Sacchi 2006; Xu et al. 2019), however, its efficiency when added to the aqueous phases of soil RNA extraction has not yet been reported. Mettel *et al*. (Mettel et al. 2010) used an acidic phenol (pH = 4.5) to separate DNA from RNA in soil samples, but our results, as well as those of a recently published study (de la Rubia et al. 2022), show that aqueous pH is more important than the pH of the added phenol solution when separating DNA from RNA (de la Rubia et al. 2022). Interestingly, we observed that the addition of low-pH sodium acetate in this step also slightly reduced the amount of humic substances (Supplementary Table 1). To compare and verify the performance of the improved method with the mentioned methods (Angel et al. 2012; Griffiths et al. 2000; Sharma et al. 2012; Thorn et al. 2019), the extraction related to these methods was also investigated in terms of quantity and quality (Fig. 4 and Table 2). RNA integrity was assessed by electrophoresis of total RNA on agarose gels. On the gel, distinct bands were observed for 23S and 16S rRNA (Fig. 4). In total, 2.8 and 2.3 μg/g of total environmental RNA were extracted from mineral and organic samples, respectively, using the different methods. Accordingly, A260/A280 and A260/A230 ratios ranged from 1.82 to 1.89 and 1.79 to 1.98, respectively (Table 2). The absorbance at A400 was also used to measure the amount of co-extracted humic substances with RNA (Mettel et al. 2010). With the improved method, the highest quality and quantity of RNA and total nucleic acid (TNA) were co-extracted with the lowest amounts of humic substances compared to the four existing methods. Additionally, a comparative extraction was performed in order to determine the effect of low-pH sodium acetate buffer on reducing the amount of humic substances. The results showed that, in the sodium acetate-free situation, not only was the amount of humic acid slightly higher but the quality of the extracted RNA was also less than other situations (Supplementary Table 1). Additionally, price comparisons revealed that this improved protocol was significantly cheaper than the commercial kit to obtain high-quality DNA/RNA from soil samples (Supplementary File S1).

**Fig. 4.**
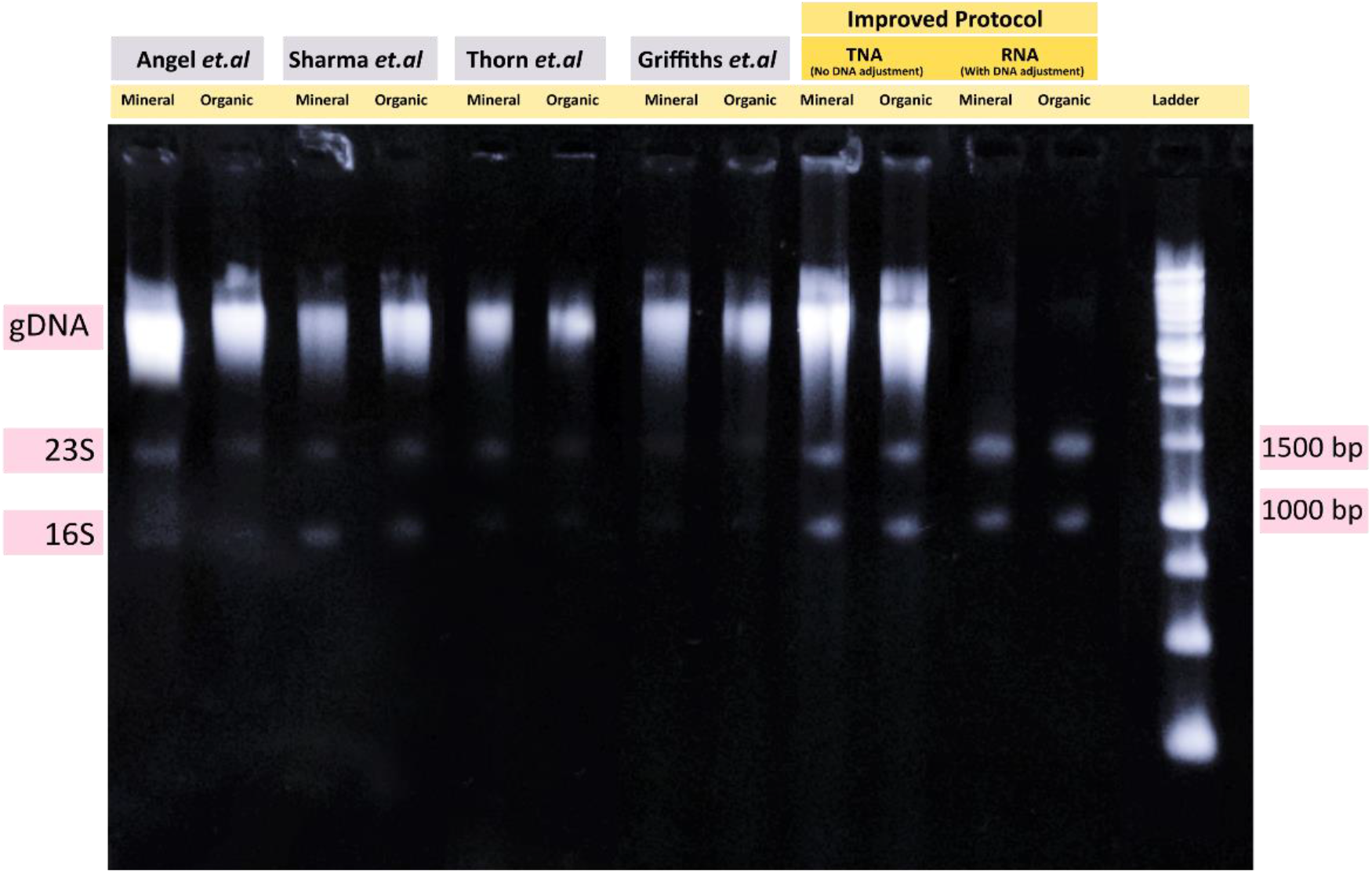
Gel electrophoresis of extracted RNA and total nucleic acid (TNA) following four existing methods (Griffiths et al. (Griffiths et al. 2000), Thorn et al. (Thorn et al. 2019), Sharma et al. (Sharma et al. 2012), and Angel et al. (Angel et al. 2012)) and the improved method from mineral and organic soil. In the four wells on the right side, total RNA extraction with DNA adjustment (using low-pH sodium acetate buffer) as well as TNA (without adjusting DNA) using the improved method loaded. The nucleic acids were visualized on 1% agarose-SB gel.

**Table 2.** Qualitative and quantitative properties of total RNA extracted from mineral and organic soil samples using five different extraction methods

| Protocol | Sample ID | Nucleic Acid (ng/μl) | 260/280 | 260/230 | A400 nm | Qubit RNA (ng/μl) | Qubit DNA (ng/μl) |
| --- | --- | --- | --- | --- | --- | --- | --- |
| Griffiths *et al.* | Organic | 24.7 | 1.51 | 0.89 | 0.09 | 2.01 | 17.1 |
|  | Mineral | 27.3 | 1.69 | 1.42 | 0.078 | 3.23 | 21 |
| Angel *et al.* | Organic | 23.4 | 1.73 | 1.12 | 0.019 | 3.02 | 16 |
|  | Mineral | 36 | 1.8 | 1.59 | 0.013 | 3.9 | 23.1 |
| Thorn *et al.* | Organic | 23 | 1.58 | 0.79 | 0.021 | 2.22 | 15.63 |
|  | Mineral | 25 | 1.79 | 1.23 | 0.018 | 2.9 | 18.2 |
| Sharma *et al.* | Organic | 27 | 1.8 | 1.58 | 0.02 | 2.1 | 19.1 |
|  | Mineral | 29 | 1.86 | 1.68 | 0.011 | 3.08 | 21.97 |
| Improved method | Organic | 56.6 | 1.82 | 1.79 | 0.006 | 9.3 | 39.8 |
|  | Mineral | 78 | 1.89 | 1.98 | 0.004 | 11.2 | 61.2 |

### Direct RNA sequencing

The primary objective of this study was to improve RNA extraction from organic and mineral soil for metatranscriptomics studies, so the nanopore direct RNA sequencing kit was chosen for library preparation. In the first step, 300 ng of each extracted RNA sample from mineral or organic soil samples was split into two groups, and four libraries with four custom barcodes were prepared and pooled together. After 24 hours, 306,953 reads with ~215 million bp were sequenced successfully. The 10.1 mean read quality score with 702 bp mean read length was achieved for the pooled library (Supplementary Table 2).

### Bioinformatics analysis

#### Preprocessing the data

First, raw data were base-called, filtered, and demultiplexed, and then the quality of the filtered reads (274,565) was assessed using the NanoPlot software (Supplementary Fig. 2). For mineral total RNA, mineral mRNA, organic total RNA, and organic mRNA, the total number of megabases was 52.2, 80.1, 48.6 and 18.1 while the number of reads was 80,219, 108,114, 59,453 and 26,779, respectively (Table 3). For mineral total RNA, mineral mRNA, organic total RNA, and organic mRNA, the N50 read lengths were 887, 1087, 1254, and 972 bp, and the maximum read lengths were 2.9, 3.6, 3.5, and 2.8 kb, respectively. Mineral mRNA and organic total RNA samples provided many long reads with a median read length of 789 and 658 bp, while mineral total RNA and organic mRNA samples only provided median read lengths of 560 and 583 bp (Supplementary File S2).

**Table 3.** Summary of direct RNA sequencing results for mineral and organic soil samples

| Sample | Trimmed<br>(n<sup>5</sup>) | Total<br>bases | Read length |  |  |
| --- | --- | --- | --- | --- | --- |
|  |  |  | Minimum | Average | Maximum |
| MTR<sup>1</sup> | 80,219 | 52,496,760 | 36 | 654 | 2945 |
| OTR<sup>2</sup> | 59,453 | 48,640,361 | 2 | 818 | 3575 |
| MMR<sup>3</sup> | 108,114 | 80,095,001 | 25 | 740 | 3602 |
| OMR<sup>4</sup> | 26,779 | 18,138,860 | 11 | 677 | 2825 |

1. Mineral total RNA; 2. Organic total RNA; 3. Mineral mRNA; 4. Organic mRNA; 5.
Number of read

### Clustering and taxonomic classification

RATTLE’s clustering and correction pipeline constructs a consensus sequence from each cluster associated with each barcode for taxonomic classification. Total RNA samples from minerals and organics contained 10.7% and 16.7% of ribosomal RNA (rRNA) clusters, respectively. The rRNA clusters were retained for taxonomic classification. Mineral and organic rRNA clusters showed that 77% and 81% of rRNAs belong to the bacteria kingdom, respectively. A large portion of the bacteria group in mineral soil comes from the Proteobacteria group (45%), while the Terrabacteria group (42%) dominates in organic soil (Fig. 5 and Supplementary File S3-4).

**Fig. 5.**
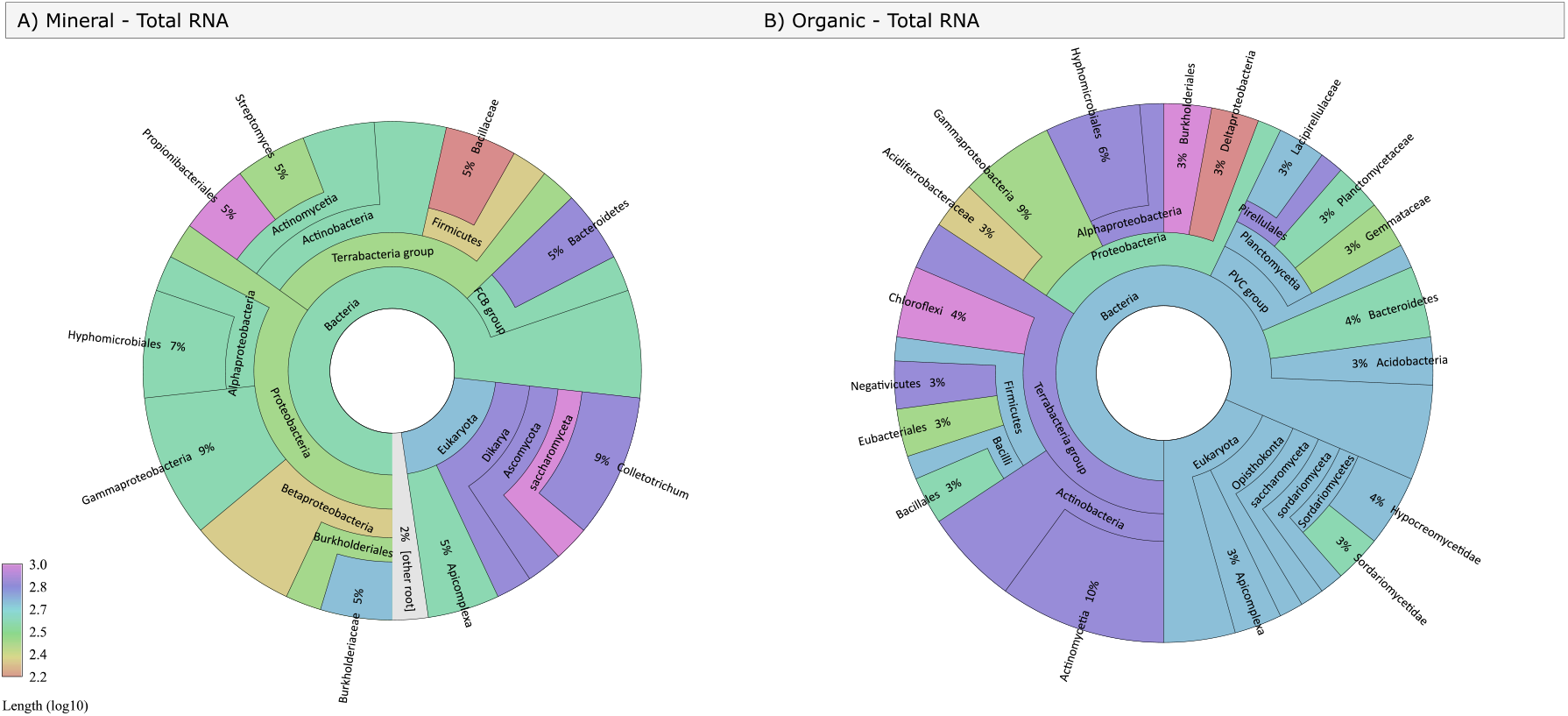
Taxonomic plots for mineral rRNA (A), organic rRNA (B). The comparative analysis and visualization of results were done by Recentrifuge. Based on its developer’s suggestion for nanopore sequencing reads, the LOGLENGTH was considered as a scoring scheme (Martí 2019). For detailed prospection of the results, interactive graphs are available in the supplementary materials (Supplementary File S3-4).

In addition, the remaining non-rRNAs, mineral mRNAs, and organic mRNAs were taxonomically categorized by Kraken 2. A nucleotide search was performed by Kraken 2 against the PlusPF database (May 2021). Kraken 2 programs detected a diversity of microorganisms, including bacteria, Archaea, and fungi in mineral and organic soils (Fig. 6). For example, in the mineral total RNA sample, seven bacterial phyla were identified, including Actinobacteria, Bacteroidota, Chloroflexota, Deinococcota, Firmicutes, Planctomycetota, and Proteobacteria. This sample also contained one fungal phylum, Ascomycota, and two Archaea phyla, Euryarchaeota and Thaumarchaeota. Furthermore, the presence of various proteobacteria, including *Pseudomonas spp*., was dominant in both mineral (Fig. 6A and 6B) and organic soil samples (Fig. 6C and 6D). However, just a single cluster was classified as belonging to the virus kingdom Orthornavirae in the mineral mRNA sample (Fig. 6B).

**Fig. 6.**
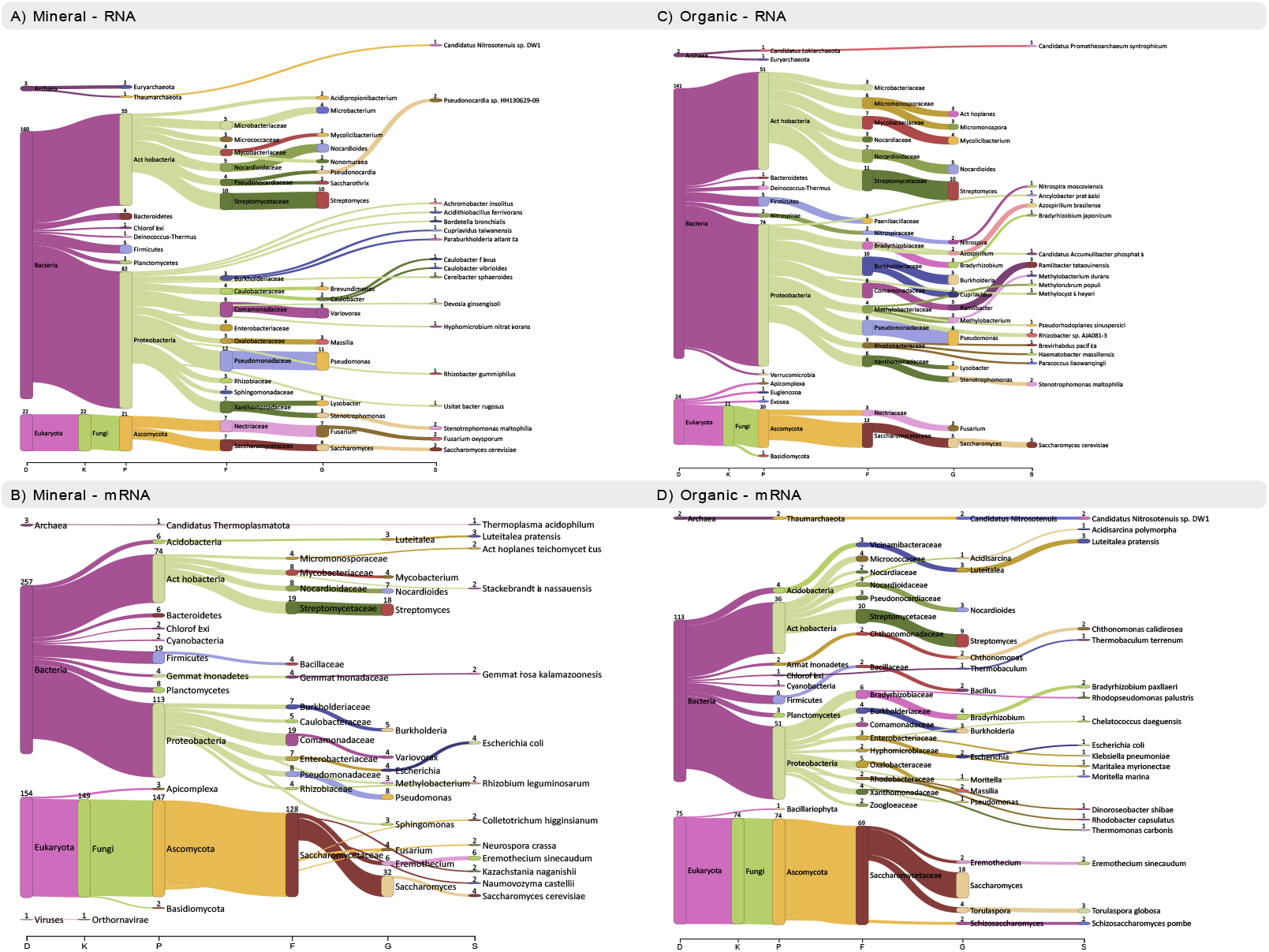
Taxonomical classification of mineral and organic soil metatranscriptomes at various levels: kingdom (K), phylum (P), class (C), order (O), family (F), and genus (G). Visualization of results was done using Pavian (Breitwieser and Salzberg 2020).

Since approximately 30% of the clusters were not aligned by Kraken 2, the unaligned clusters were searched against a custom database using Centrifuge. *Acanthamoeba castellanii*, a free-living soil amoeba, was detected in unaligned clusters of mineral samples (both total RNA and mRNA) (Supplementary Figs. 3A and 3B). The centrifuge analysis revealed that Fusarium genus was also present in mineral and organic non-hit mRNA clusters (Supplementary Figs. 3B and 3D).

### Annotation and functional analysis

As a first step in evaluating and interpreting the content of our metatranscriptomes, we blasted the clusters and identified ORFs using BLASTx and BLASTp, respectively, against the SwissProt database with an E-value cut-off set to 10^-5^. Clusters of Orthologous Groups (COGs) analyses were performed on both mineral and organic read clusters. In both mineral and organic samples, COGs related to translation, ribosomal structure, and biogenesis had the highest cluster abundance. Flagellar biosynthesis-related clusters in organic samples were less expressed than other clusters. Meanwhile, the lowest level of COGs in mineral cluster was associated with different functions, including defense mechanisms, intracellular trafficking, secretion, and vesicular transport (Fig. 7).

**Fig. 7.**
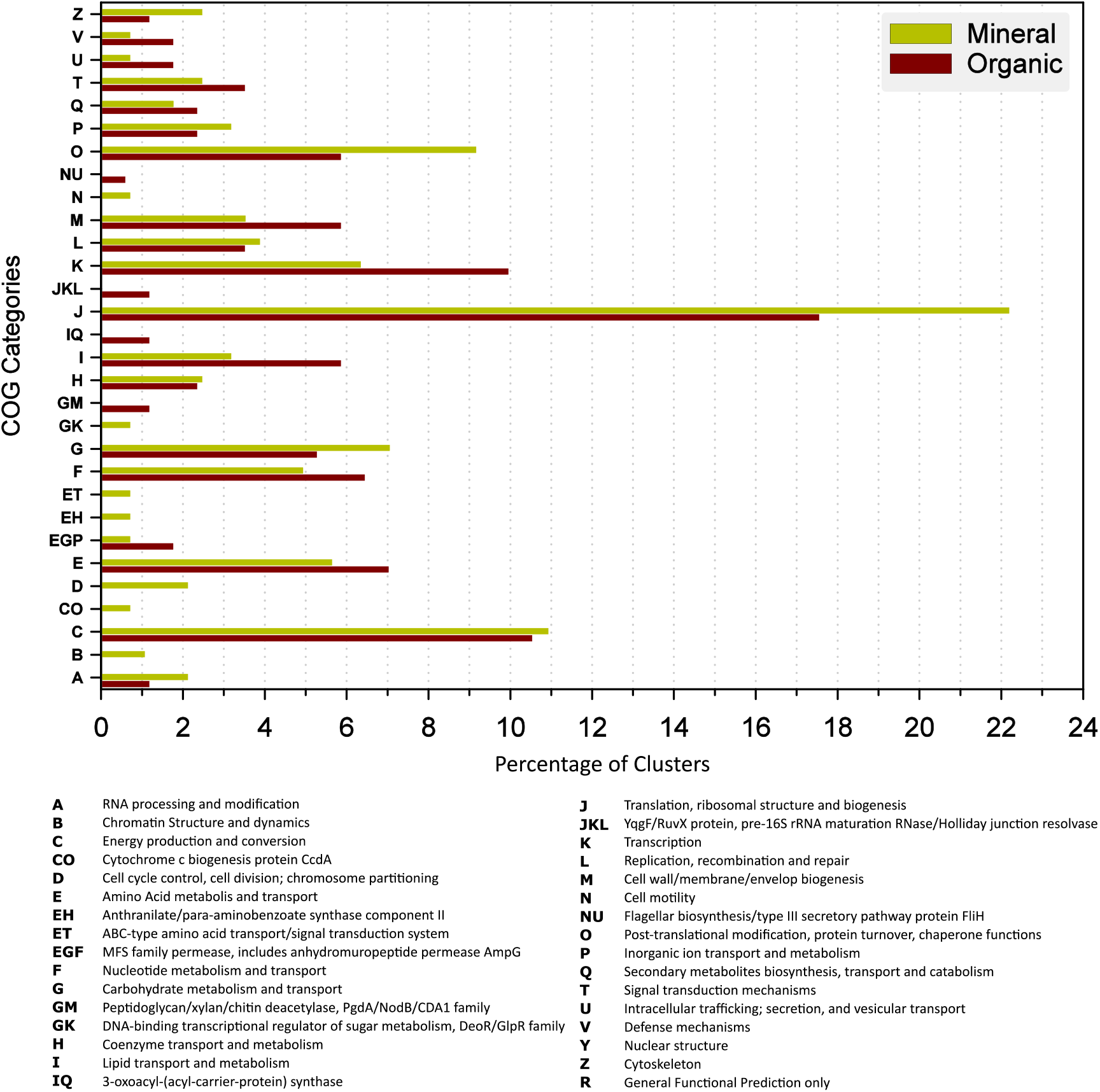
Functional classification of Cluster of Orthologous Groups (COG) on mineral and organic clusters. Classification extracted from eggNOG-mapper2 functional analysis (Cantalapiedra et al. 2021). 336 mineral clusters and 221 organic clusters were assigned to 30 COG categories. Functional classes reflect specific genes and metabolisms, as well as environmental factors. The results were visualized by Sigmaplot v14.5 (Systat Software, San Jose, CA) and Inkscape (https://inkscape.org).

To achieve the community’s functioning, gene ontology (GO) classification was taken into account using SwissProt protein sequences. GO terms are categorized as biological processes, cellular components, and molecular functions, which describe the properties of a gene product (Ashburner et al. 2000; Gene Ontology 2004). It was observed that the relative distribution of GO terms across mineral and organic datasets varied, indicating that each mineral and organic sample was inhabited by a different metabolically active community of microorganisms (8).

In mineral and organic soil, gene ontology analysis assigned 495 and 288 clusters, respectively, to one or more GO terms. The three most common categories of biological process for mineral soils and organic soils were “metabolic process (GO:0008152)” with 120 and 74 clusters, “nitrogen compound metabolic process (GO:0006807)” with 102 and 65 clusters, and “biosynthetic process (GO:0009058)” with 73 and 45 clusters, respectively. In the category of cellular components, the three enriched terms with the most clusters in the mineral soil sample were cytoplasm (GO:0005737), membrane (GO:0016020), and ribosome (GO:0005840). Among enriched terms within the cellular component category in the organic soil sample, “cytoplasm (GO:0005737)” ranked first with 28 clusters, followed by “membrane (GO:0016020)” ranking second with 25 clusters, and “plasma membrane (GO:0005886)” ranking third with 17 clusters. The three most abundant groups in the molecular function domain for the mineral soil sample were “catalytic activity (GO:0003824)” with 104 clusters, “ion binding (GO:0043167)” with 91 clusters, and “DNA-binding (GO:0003676)” with 65 clusters. For the molecular function domain in the organic soil sample, catalytic activity (GO:0003824) was the most abundant group, with 74 clusters, followed by ions binding (GO:0043167), with 57 clusters, and metal ion binding (GO:0046872), with 38 clusters (Fig. 8 and Supplementary File S5-6).

**Fig. 8.**
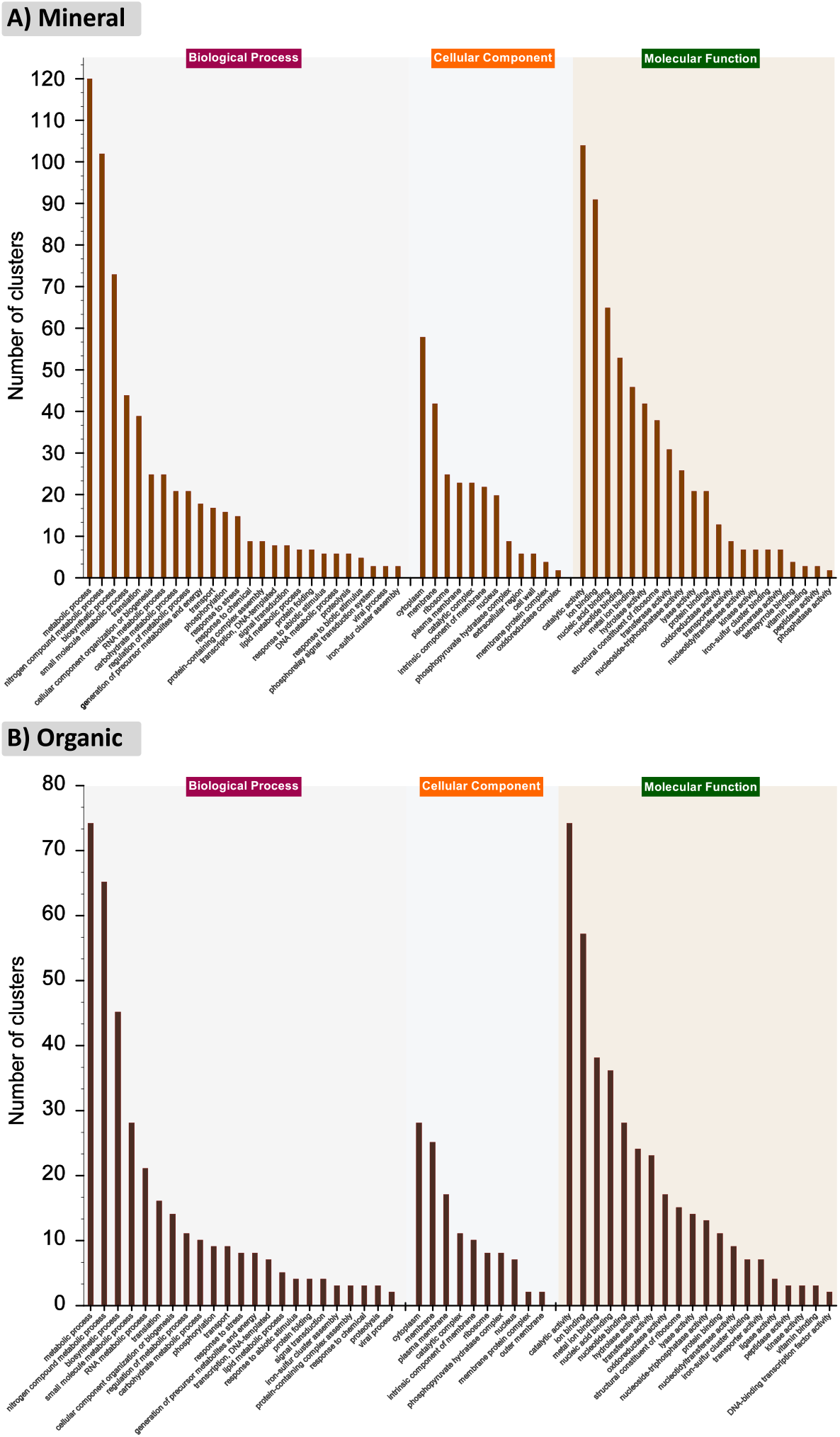
Gene ontology enrichment analysis of mineral and organic clusters. There are three categories of gene ontology terms for each mineral (A) and organic (B) cluster: biological processes, cellular components, and molecular functions. The results were visualized by Sigmaplot v14.5 (Systat Software, San Jose, CA) and Inkscape (https://inkscape.org).

A total of 359 and 261 numbers of mineral and organic KOs were classified, respectively. The top three most abundant categories at the BRITE 1 functional hierarchy level were related to “environmental information processing”, “metabolism”, and “genetic information processing” with 36.9%, 32.8% and 19.9% of organic KOs and 33%, 29.3% and 17.9% of mineral KOs, respectively. However, a comparison between mineral and organic KOs demonstrated that the cellular progress category had greater activation in the mineral samples (Fig. 9B).

**Fig. 9.**
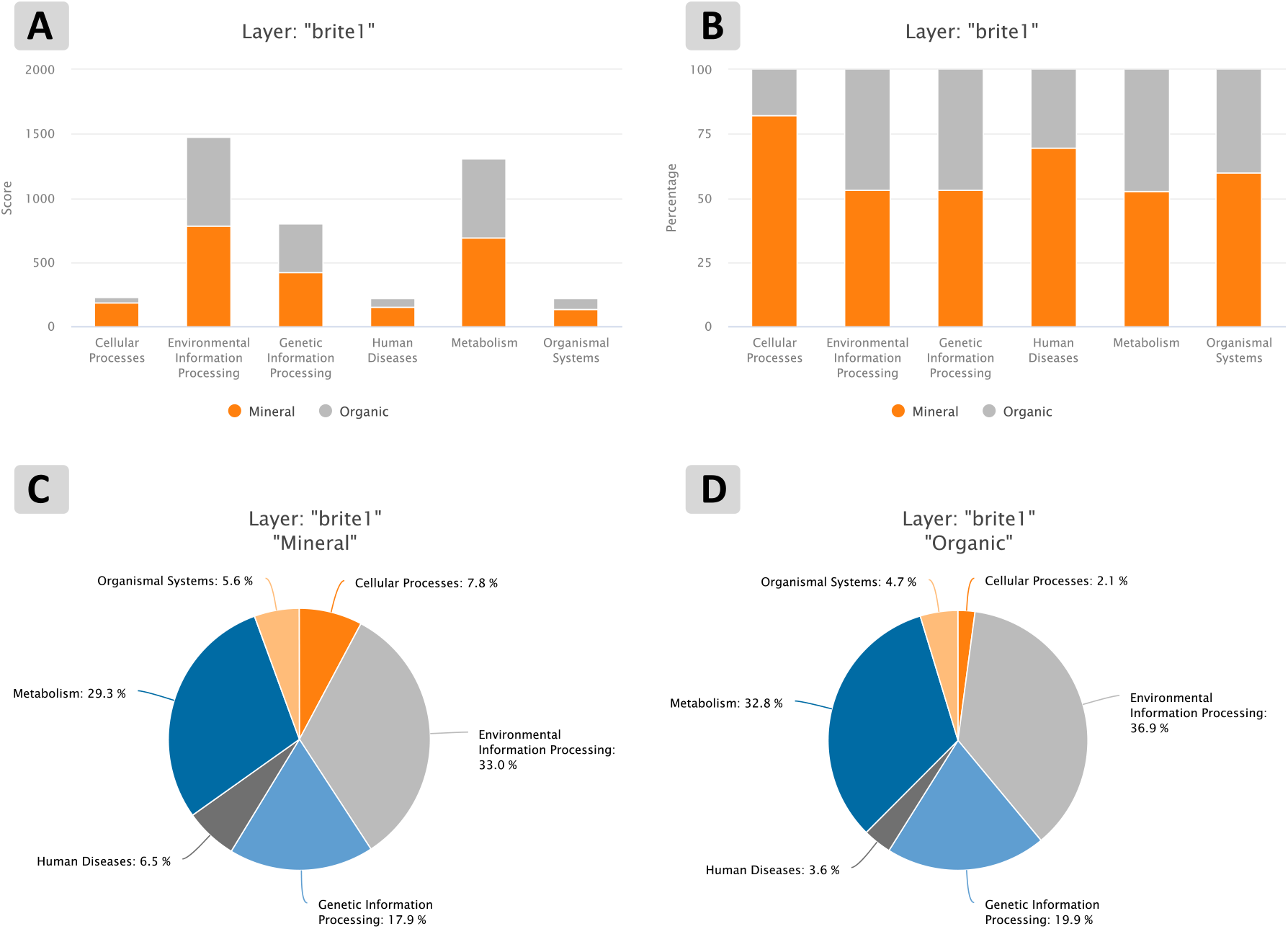
KEEG BRITE 1 analysis. Stacked barplots and pie charts of mineral and soil clusters involved in KEGG BRITE 1. (A and B) Stacked barplots of mineral and organic data series before and after the normalization step. (C and D) Pie charts of mineral and organic clusters

At the BRITE 2 functional hierarchy level, mineral and organic samples showed several significant differences. The three most abundant categories in the organic samples at the BRITE 2 level were “Drug resistance: Antimicrobial”, “Development”, and “Digestive systems” which were not present in the mineral sample (Supplementary Fig. 4B). In the mineral sample, the most abundant categories at the BRITE 2 functional hierarchy level were “Drug resistance: Antineoplastic”, and “Immune diseases” which were absent in the organic sample. Nevertheless, the third-highest category was “Cellular community: eukaryotes” which was also activated slightly in the organic sample (Supplementary Fig. 4B).

By using FuncTree 2 (Darzi et al. 2019), functional hierarchies were also visualized at different levels of the BRITE functional hierarchy (BRITE 1, BRITE 2) and pathway (Supplementary Fig. 5A). A total of 41 and 42 functional categories from the BRITE 2 level, which were assigned to six categories at the BRITE 1 level, were activated in the mineral and organic samples, respectively. A total of 176 and 159 pathways were enriched from mineral and organic clusters, respectively, and 99 pathways were the same between mineral and organic soil (Supplementary File S7). For instance, lipid metabolism pathways were enriched specifically in the organic sample (Supplementary Fig. 6A), while in the mineral sample carbohydrate metabolism pathways were particularly active (Supplementary Fig. 6B).

## Discussion

Various methods have been proposed over the last three decades for extracting nucleic acids of soil microbial populations, and with time, these methods have been refined and improved. Aside from enhancing efficiency and being responsive to current technology requirements (e.g., quantity and quality), these changes also aimed to address the primary challenges associated with nucleic acid extraction from soil, such as co-extraction of contaminants, nucleic acid absorption into soil particles, and use of adequate pH. The nature of RNA and its instability make extracting RNA from soil more difficult, along with other soil challenges. Consequently, we labored to improve several existing methods (Angel et al. 2012; Griffiths et al. 2000; Peršoh et al. 2008; Sharma et al. 2012; Thorn et al. 2019) with modifications based on previous research (Guerra et al. 2020; Hashizume 2015; Mettel et al. 2010; Wang et al. 2009a; Wang et al. 2009b) to achieve a novel, and cost-effective RNA extraction method that performed strikingly well in mineral and organic soils.

The method presented was designed to reach two key goals: to (1) prevent the addition of alkaline pH to the nucleic acid extraction; and (2) prevent the co-extraction of humic acids with the nucleic acid by performing all extractions on ice. Indeed, processing samples at a low temperature substantially reduces humic acid carryover, protects RNA from degradation, and prevents tubes from being accidentally overheated and leaking phenol (Angel et al. 2012). Here, a CTAB buffer containing PVP (Sharma et al. 2012) was modified to reach a lower pH. However, it should be noted that while lower pH prevents humic compounds from being extracted, it also reduces the concentration of nucleic acid, and we observed that pHs below 5.5 significantly reduced nucleic acid concentration. The use of phosphate buffers separately for extraction has two benefits: modification of pH levels (according to soil pH and buffer pH) and an increase in phosphate content in the lysis step, which helps to release nucleic acids from soil particles (Guerra et al. 2020). Depending on the soil type and pH of the phenol solution used, this step of the protocol can be adjusted in clay-rich soils by increasing the phosphate molarity (above 150 mM) without changing the CTAB buffer. On the other hand, if the available phenol had a pH that was above 6 or below 5, the extraction buffer pH can be modified by changing the pH of the phosphate buffer, which is much safer and simpler than changing the pH of the phenol. We tested a variety of phenols, including citrate-phenol, Tris-saturated phenol, Trizol, and water-saturated phenol with pH 5 and 4 in our pre-experiments, which led us to choose water-saturated phenol with pH 5.8 based on the yield and purity of nucleic acids (data not shown). Low-quality results with the use of these different phenol solutions can be attributed to their salt content (Tris, Citrate, Guanidinium thiocyanate, etc.). Through the addition of the phenol-chloroform step and re-purification of the aqueous phase, we can adjust the amount of DNA using acidic sodium acetate buffer (Chomczynski and Sacchi 2006).

Additionally, our results revealed that sodium acetate could absorb and reduce humic contamination to some extent. Although sodium acetate has been used previously to absorb humic substances (Lafrance and Mazet 1989; Rashid 1969), we found that low-pH sodium acetate helped to decrease humic substances as well as to separate DNA from RNA. However, the effect of sodium acetate on humic acid reduction will need further investigation, because this type of contamination is common in different soil types and different ecosystems (Supplementary Table 1). Contamination is a very critical issue in a lot of molecular biology techniques (e.g., PCR, RT-PCR, macromolecule blotting and probing, sequencing), especially when working with nanopore direct RNA kit, because the RNA must be contamination-free, and the enzymatic steps such as polyadenylation, ligation, and reverse transcription are extremely sensitive.

Indeed, several challenges remain when analyzing nanopore data, and the initial stages are still among the most challenging. This is striking when it comes to working with direct RNA sequencing data since the development of dedicated and user-friendly bioinformatics tools for this purpose is ongoing. To answer this need, we designed in this study an in-house bioinformatics workflow for taxonomy analysis at different levels. Notably, using the RATTLE Pipeline (de la Rubia et al. 2022), it was possible to avoid misclassifying the reads of species with the least sequencing coverage, one of the common issues when working with long-read assemblers. Thus, using this in-house workflow we analyzed the consensus sequences obtained from the clustering step in the total RNA datasets in terms of rRNA sequences and we completed the taxonomic profiling of the microbial population.

The remaining reads were analyzed by Kraken2 and Centrifuge tools (Kim et al. 2016; Wood et al. 2019). The noteworthy point in this context is Kraken2’s insufficient ability to fully classify sequences. However, due to its much higher speed compared to Centrifuge and updated databases, it can still be useful as an early step to obtain the microbiome profile. On the other hand, the Centrifuge needs an updated or dedicated database for the desired project and also requires more time and system resources. Nevertheless, the use of both tools together enabled the loss of sequences without hits to be minimized. The identification of Orthornavirae viruses in the mRNA dataset demonstrates the high performance of the RNA enrichment method to capture mRNA from low-relative-abundance microbial species (Fig. 6B), especially for viruses that have polyA-tailed mRNA (Brinton et al. 2021; Lang et al. 2021; Walker et al. 2021).

A major advantage of using RNA datasets is that it allows for functional analysis and annotation to be performed. The importance of this issue becomes even more apparent when dealing with soil environments, because it allows for determining the active functional profile in the microbiome (which cannot always be determined by examining DNA) and, ultimately, better understanding the plant-soil interaction. In addition to the difference in their microbial profile the annotation results (GO, KEGG, and COG analysis) showed a higher functional activity in the mineral soil than in the organic one. However, this difference in functional activity maybe driven by the number of clusters and reads obtained from these two types of soil. Indeed, organic soil also showed greater activity in “Cell wall/membrane/envelope biogenesis”, “Lipid transport and metabolism”, and “Transcription” groups (Fig. 7). Therefore, RNA nanopore sequencing results should be combined with short reads sequencing results of the same samples to obtain a suitable coverage for accurate functional analysis of genes with low expression frequency.

## Conclusion

Several of the challenges associated with soil nucleic acid extraction have been addressed by the development of soil nucleic acid extraction kits. However, due to the high cost of kits for projects that require a large number of samples, as well as the impossibility of optimizing these kits for different types of soils, manual methods have continued to be of great interest. We present a new improved method to extract RNA from soil, which combines the strengths of the previously developed methods, but also adds steps to adjust the amount of DNA and increase the quality and quantity of extracted material in two soil types (mineral and organic). As a result of using nanopore direct RNA sequencing, it was possible to confirm the high quality and purity of the final product. Furthermore, an optimal workflow for sequencing data clustering, taxonomy analysis, and functional annotation has been proposed and evaluated. To our knowledge, this work breaks ground by sequencing soil RNA with direct RNA nanopore sequencing and most conveniently proposes an in-house bioinformatics workflow necessary to process this type of dataset. In this third-generation sequencing era, the yield, quality and purity of the RNA obtain from this improved cost-effective method open the door to decipher the soil microbial functionalities using nanopore sequencing technology. Moreover, we proposed a strong and straightforward bioinformatics workflow to analysis the nanopore sequencing data resulting from this improved RNA extraction method.

## Supporting information

Supplementary HTML Graphs

## Acknowledgment

The authors gratefully acknowledge Pierre Lemoyne, Azza Larafa and Dong Xu from Agriculture and Agri-Food Canada’s Saint-Jean-sur-Richelieu Research and Development Centre (CRDH) for their support and help on soil sampling and processing. The authors would like to thank Dave Thibouthot-Ste-Croix, from the nematology lab. of the CRDH, for his technical support on nanopore sequencing and Joel Lafond-Lapalme, our team leader in bioinformatics, for his assistance for finding and resolving bugs.

## Literature cited

Aguiar-Pulido, V., Huang, W., Suarez-Ulloa, V., Cickovski, T., Mathee, K., and Narasimhan, G. 2016. Metagenomics, Metatranscriptomics, and Metabolomics Approaches for Microbiome Analysis: Supplementary Issue: Bioinformatics Methods and Applications for Big Metagenomics Data. Evolutionary Bioinformatics 12s1:EBO.S36436.

Alm Elizabeth, W., Zheng, D., and Raskin, L. 2000. The Presence of Humic Substances and DNA in RNA Extracts Affects Hybridization Results. Applied and Environmental Microbiology 66:4547–4554.

Altschul, S. F., Gish, W., Miller, W., Myers, E. W., and Lipman, D. J. 1990. Basic local alignment search tool. Journal of Molecular Biology 215:403–410.

Angel, R., Claus, P., and Conrad, R. 2012. Methanogenic archaea are globally ubiquitous in aerated soils and become active under wet anoxic conditions. The ISME Journal 6:847–862.

Ashburner, M., Ball, C. A., Blake, J. A., Botstein, D., Butler, H., Cherry, J. M., Davis, A. P., Dolinski, K., Dwight, S. S., Eppig, J. T., Harris, M. A., Hill, D. P., Issel-Tarver, L., Kasarskis, A., Lewis, S., Matese, J. C., Richardson, J. E., Ringwald, M., Rubin, G. M., and Sherlock, G. 2000. Gene Ontology: tool for the unification of biology. Nature Genetics 25:25–29.

Azeem, M., Soundari, P. G., Ali, A., Tahir, M. I., Imran, M., Bashir, S., Irfan, M., Li, G., Zhu, Y.-G., and Zhang, Z. 2021. Soil metaphenomics: a step forward in metagenomics. Archives of Agronomy and Soil Science:1–19.

Bairoch, A., and Apweiler, R. 1997. The SWISS-PROT protein sequence data bank and its supplement TrEMBL. Nucleic Acids Research 25:31–36.

Breitwieser, F. P., and Salzberg, S. L. 2020. Pavian: interactive analysis of metagenomics data for microbiome studies and pathogen identification. Bioinformatics 36:1303–1304.

Brinton, M. A., Gulyaeva, A. A., Balasuriya, U. B. R., Dunowska, M., Faaberg, K. S., Goldberg, T., Leung, F. C. C., Nauwynck, H. J., Snijder, E. J., and Stadejek, T. 2021. ICTV virus taxonomy profile: Arteriviridae 2021. The Journal of General Virology 102.

Cantalapiedra, C. P., Hernández-Plaza, A., Letunic, I., Bork, P., and Huerta-Cepas, J. 2021. eggNOG-mapper v2: Functional Annotation, Orthology Assignments, and Domain Prediction at the Metagenomic Scale. Molecular Biology and Evolution 38:5825–5829.

Cavicchioli, R., Ripple, W. J., Timmis, K. N., Azam, F., Bakken, L. R., Baylis, M., Behrenfeld, M. J., Boetius, A., Boyd, P. W., Classen, A. T., Crowther, T. W., Danovaro, R., Foreman, C. M., Huisman, J., Hutchins, D. A., Jansson, J. K., Karl, D. M., Koskella, B., Mark Welch, D. B., Martiny, J. B. H., Moran, M. A., Orphan, V. J., Reay, D. S., Remais, J. V., Rich, V. I., Singh, B. K., Stein, L. Y., Stewart, F. J., Sullivan, M. B., van Oppen, M. J. H., Weaver, S. C., Webb, E. A., and Webster, N. S. 2019. Scientists’ warning to humanity: microorganisms and climate change. Nature Reviews Microbiology 17:569–586.

Chaparro-Encinas, L. A., Arellano-Wattenbarger, G. L., Parra-Cota, F. I., and de los Santos-Villalobos, S. 2020. A modified CTAB and Trizol^®^ protocol for high-quality RNA extraction from whole wheat seedlings, including rhizosphere. Cereal Research Communications 48:275–282.

Chomczynski, P., and Sacchi, N. 2006. The single-step method of RNA isolation by acid guanidinium thiocyanate–phenol–chloroform extraction: twenty-something years on. Nature Protocols 1:581–585.

Darzi, Y., Yamate, Y., and Yamada, T. 2019. FuncTree2: an interactive radial tree for functional hierarchies and omics data visualization. Bioinformatics 35:4519–4521.

De Coster, W., D’Hert, S., Schultz, D. T., Cruts, M., and Van Broeckhoven, C. 2018. NanoPack: visualizing and processing long-read sequencing data. Bioinformatics 34:2666–2669.

de la Rubia, I., Srivastava, A., Xue, W., Indi, J. A., Carbonell-Sala, S., Lagarde, J., Albà, M. M., and Eyras, E. 2022. RATTLE: Reference-free reconstruction and quantification of transcriptomes from Nanopore sequencing. bioRxiv:2020.2002.2008.939942.

Deutscher, M. P. 2006. Degradation of RNA in bacteria: comparison of mRNA and stable RNA. Nucleic Acids Research 34:659–666.

Fischer, M., Renevey, N., Thür, B., Hoffmann, D., Beer, M., and Hoffmann, B. 2016. Efficacy Assessment of Nucleic Acid Decontamination Reagents Used in Molecular Diagnostic Laboratories. PLOS ONE 11:e0159274.

Gene Ontology, C. 2004. The Gene Ontology (GO) database and informatics resource. Nucleic Acids Research 32:D258–D261.

Goring, C. A. I., and Bartholomew, W. V. 1952. ADSORPTION OF MONONUCLEOTIDES, NUCLEIC ACIDS, AND NUCLEOPROTEINS BY CLAYS. Soil Science 74.

Griffiths, R. I., Whiteley, A. S., O’Donnell, A. G., and Bailey, M. J. 2000. Rapid method for coextraction of DNA and RNA from natural environments for analysis of ribosomal DNA-and rRNA-based microbial community composition. Applied and Environmental Microbiology 66:5488–5491.

Guerra, V., Beule, L., Lehtsaar, E., Liao, H.-L., and Karlovsky, P. 2020. Improved Protocol for DNA Extraction from Subsoils Using Phosphate Lysis Buffer. Microorganisms 8.

Hashizume, H. 2015. Adsorption of Nucleic Acid Bases, Ribose, and Phosphate by Some Clay Minerals. Life 5.

Hayden, H. L., Savin, K. W., Wadeson, J., Gupta, V. V. S. R., and Mele, P. M. 2018. Comparative Metatranscriptomics of Wheat Rhizosphere Microbiomes in Disease Suppressive and Non-suppressive Soils for Rhizoctonia solani AG8. Frontiers in Microbiology 9.

Huerta-Cepas, J., Szklarczyk, D., Heller, D., Hernández-Plaza, A., Forslund, S. K., Cook, H., Mende, D. R., Letunic, I., Rattei, T., Jensen, Lars J., von Mering, C., and Bork, P. 2019. eggNOG 5.0: a hierarchical, functionally and phylogenetically annotated orthology resource based on 5090 organisms and 2502 viruses. Nucleic Acids Research 47:D309–D314.

Jansson, J. K., and Hofmockel, K. S. 2020. Soil microbiomes and climate change. Nature Reviews Microbiology 18:35–46.

Kanehisa, M., and Goto, S. 2000. KEGG: Kyoto Encyclopedia of Genes and Genomes. Nucleic Acids Research 28:27–30.

Kim, D., Song, L., Breitwieser, F. P., and Salzberg, S. L. 2016. Centrifuge: rapid and sensitive classification of metagenomic sequences. Genome Research 26:1721–1729.

Kopylova, E., Noé, L., and Touzet, H. 2012. SortMeRNA: fast and accurate filtering of ribosomal RNAs in metatranscriptomic data. Bioinformatics 28:3211–3217.

Lafrance, P., and Mazet, M. 1989. Adsorption of Humic Substances in the Presence of Sodium Salts. Journal AWWA 81:155–162.

Lang, A. S., Vlok, M., Culley, A. I., Suttle, C. A., Takao, Y., Tomaru, Y., and Consortium, I. R. 2021. ICTV virus taxonomy profile: Marnaviridae 2021. The Journal of General Virology 102.

Lever, M. A., Torti, A., Eickenbusch, P., Michaud, A. B., Šantl-Temkiv, T., and Jørgensen, B. B. 2015. A modular method for the extraction of DNA and RNA, and the separation of DNA pools from diverse environmental sample types. Frontiers in Microbiology 6.

Lim, N. Y. N., Roco, C. A., and Frostegård, Å. 2016. Transparent DNA/RNA Co-extraction Workflow Protocol Suitable for Inhibitor-Rich Environmental Samples That Focuses on Complete DNA Removal for Transcriptomic Analyses. Frontiers in Microbiology 7.

Martí, J. M. 2019. Recentrifuge: Robust comparative analysis and contamination removal for metagenomics. PLOS Computational Biology 15:e1006967.

Mehlich, A. 1984. Mehlich 3 soil test extractant: A modification of Mehlich 2 extractant. Communications in Soil Science and Plant Analysis 15:1409–1416.

Mettel, C., Kim, Y., Shrestha Pravin, M., and Liesack, W. 2010. Extraction of mRNA from Soil. Applied and Environmental Microbiology 76:5995–6000.

Paulin, M. M., Nicolaisen, M. H., Jacobsen, C. S., Gimsing, A. L., Sørensen, J., and Bælum, J. 2013. Improving Griffith’s protocol for co-extraction of microbial DNA and RNA in adsorptive soils. Soil Biology and Biochemistry 63:37–49.

Pei, Y., Mamtimin, T., Ji, J., Khan, A., Kakade, A., Zhou, T., Yu, Z., Zain, H., Yang, W., Ling, Z., Zhang, W., Zhang, Y., and Li, X. 2021. The guanidine thiocyanate-high EDTA method for total microbial RNA extraction from severely heavy metal-contaminated soils. Microbial Biotechnology 14:465–478.

Peršoh, D., Theuerl, S., Buscot, F., and Rambold, G. 2008. Towards a universally adaptable method for quantitative extraction of high-purity nucleic acids from soil. Journal of Microbiological Methods 75:19–24.

Podolyan, A., and Grelet, G.-A. 2021. Suitability of six extraction methods for isolating a large quantity of high-quality RNA from New Zealand free-draining stony soil. New Zealand Journal of Agricultural Research 64:565–575.

Poursalavati, A., Rashidi-Monfared, S., and Ebrahimi, A. 2021. Toward understanding of the methoxylated flavonoid biosynthesis pathway in Dracocephalum kotschyi Boiss. Scientific Reports 11:19549.

Qin, H., Chen, X., Tang, Y., Hou, H., Sheng, R., and Shen, J. 2016. Modified method for the extraction of mRNA from paddy soils. Biotechnology Letters 38:2163–2167.

Quast, C., Pruesse, E., Yilmaz, P., Gerken, J., Schweer, T., Yarza, P., Peplies, J., and Glöckner, F. O. 2013. The SILVA ribosomal RNA gene database project: improved data processing and web-based tools. Nucleic Acids Research 41:D590–D596.

Rajarapu, S. P., Shreve, J. T., Bhide, K. P., Thimmapuram, J., and Scharf, M. E. 2015. Metatranscriptomic profiles of Eastern subterranean termites, Reticulitermes flavipes (Kollar) fed on second generation feedstocks. BMC Genomics 16:332.

Ranjan, K., Bharti, M. K., Siddique, R. A., and Singh, J. 2021. Metatranscriptomics in Microbiome Study: A Comprehensive Approach. Pages 1–36 in: Microbial Metatranscriptomics Belowground. M. Nath, D. Bhatt, P. Bhargava and D. K. Choudhary, eds. Springer Singapore, Singapore.

Rashid, M. 1969. Contribution of Humic Substances to the Cation Exchange Capacity of Different Marine Sediments. Atlantic Geology 5:44–50.

Shakya, M., Lo, C.-C., and Chain, P. S. G. 2019. Advances and Challenges in Metatranscriptomic Analysis. Frontiers in Genetics 10.

Sharma, S., Mehta, R., Gupta, R., and Schloter, M. 2012. Improved protocol for the extraction of bacterial mRNA from soils. Journal of Microbiological Methods 91:62–64.

Sharuddin, S. S., Ramli, N., Yusoff, M. Z., Muhammad, N. A., Ho, L. S., and Maeda, T. 2022. Advancement of Metatranscriptomics towards Productive Agriculture and Sustainable Environment: A Review. International Journal of Molecular Sciences 23.

Shen, W., Le, S., Li, Y., and Hu, F. 2016. SeqKit: A Cross-Platform and Ultrafast Toolkit for FASTA/Q File Manipulation. PLOS ONE 11:e0163962.

Smith, M. A., Ersavas, T., Ferguson, J. M., Liu, H., Lucas, M. C., Begik, O., Bojarski, L., Barton, K., and Novoa, E. M. 2020. Molecular barcoding of native RNAs using nanopore sequencing and deep learning. Genome Research 30:1345–1353.

Steglich, C., Lindell, D., Futschik, M., Rector, T., Steen, R., and Chisholm, S. W. 2010. Short RNA half-lives in the slow-growing marine cyanobacterium Prochlorococcus. Genome Biology 11:R54.

Tatusov, R. L., Galperin, M. Y., Natale, D. A., and Koonin, E. V. 2000. The COG database: a tool for genome-scale analysis of protein functions and evolution. Nucleic Acids Research 28:33–36.

Thorn, C. E., Bergesch, C., Joyce, A., Sambrano, G., McDonnell, K., Brennan, F., Heyer, R., Benndorf, D., and Abram, F. 2019. A robust, cost-effective method for DNA, RNA and protein co-extraction from soil, other complex microbiomes and pure cultures. Molecular Ecology Resources 19:439–455.

Walker, P. J., Cowley, J. A., Dong, X., Huang, J., Moody, N., Ziebuhr, J., and Consortium, I. R. J. T. J. o. G. V. 2021. ICTV Virus Taxonomy Profile: Roniviridae. 102.

Wang, Y., and Fujii, T. 2011. Evaluation of Methods of Determining Humic Acids in Nucleic Acid Samples for Molecular Biological Analysis. Bioscience, Biotechnology, and Biochemistry 75:355–357.

Wang, Y., Hayatsu, M., and Fujii, T. 2009a. Extraction of bacterial RNA from soil: challenges and solutions. Microbes and environments:1202170350–1202170350.

Wang, Y., Hayatsu, M., and Fujii, T. 2012a. Extraction of Bacterial RNA from Soil: Challenges and Solutions. Microbes and Environments 27:111–121.

Wang, Y., Morimoto, S., Ogawa, N., Oomori, T., and Fujii, T. 2009b. An improved method to extract RNA from soil with efficient removal of humic acids. Journal of Applied microbiology 107:1168–1177.

Wang, Y., Nagaoka, K., Hayatsu, M., Sakai, Y., Tago, K., Asakawa, S., and Fujii, T. 2012b. A novel method for RNA extraction from Andosols using casein and its application to amoA gene expression study in soil. Applied Microbiology and Biotechnology 96:793–802.

Wang, Z., Gerstein, M., and Snyder, M. 2009c. RNA-Seq: a revolutionary tool for transcriptomics. Nature Reviews Genetics 10:57–63.

Wilson, I. G. 1997. Inhibition and facilitation of nucleic acid amplification. Applied and Environmental Microbiology 63:3741–3751.

Wnuk, E., Waśko, A., Walkiewicz, A., Bartmiński, P., Bejger, R., Mielnik, L., and Bieganowski, A. 2020. The effects of humic substances on DNA isolation from soils. PeerJ 8:e9378.

Wood, D. E., Lu, J., and Langmead, B. 2019. Improved metagenomic analysis with Kraken 2. Genome Biology 20:257.

Xu, L., Sun, L., Guan, G., Huang, Q., Lv, J., Yan, L., Ling, L., and Zhang, Y. 2019. The effects of pH and salts on nucleic acid partitioning during phenol extraction. Nucleosides, Nucleotides & Nucleic Acids 38:305–320.

Zipper, H., Buta, C., Lämmle, K., Brunner, H., Bernhagen, J., and Vitzthum, F. 2003. Mechanisms underlying the impact of humic acids on DNA quantification by SYBR Green I and consequences for the analysis of soils and aquatic sediments. Nucleic Acids Research 31:e39–e39.

Zuñiga, C., Zaramela, L., and Zengler, K. 2017. Elucidation of complexity and prediction of interactions in microbial communities. Microbial Biotechnology 10:1500–1522.

