## Supplementary Figs and Tables for "Soil metatranscriptomics: An improved RNA extraction method toward functional analysis using nanopore direct RNA sequencing"

Supplementary Table 1. Using sodium acetate at different pH levels to reduce humic acid

| **Sample ID** | **Nucleic Acid (ng/µl)** | **260/280** | **260/230** | **Abs (300nm)** | **Abs (340nm)** | **Abs (400nm)** |
|---|---|---|---|---|---|---|
| No-NaAc | 59,4 | 1,88 | 1,72 | 0,018 | 0,008 | 0,014 |
| With NaAc 3M (pH 5) | 73,6 | 1,91 | 1,83 | 0,009 | 0,001 | 0,004 |
| With NaAc 3M (pH 4.6) | 58,6 | 1,90 | 1,80 | 0,010 | 0,002 | 0,004 |
| With NaAc-HAc 2M (pH 3.4) | 35,4 | 1,93 | 1,82 | 0,012 | 0,002 | 0,002 |

Supplementary Table 2. Preliminary sequencing results for the pooled library

| | **Pooled library before trimming** | **Pooled library after trimming** |
|---|---|---|
| Mean read length | 702.5 | 717.2 |
| Median read length | 599 | 626 |
| Mean read quality | 10.1 | 11.2 |
| Median read quality | 10.4 | 11.4 |
| Number of reads | 306,953 | 285,022 |
| Read length N50 | 1,036 | 1,045 |
| Total bases | 215,635,082 | 204,427,451 |
| **Longest reads:** | | |
| 1 | 57578 | 3602 |
| 2 | 57375 | 3597 |
| 3 | 51789 | 3575 |
| 4 | 36570 | 3543 |
| 5 | 35236 | 3539 |

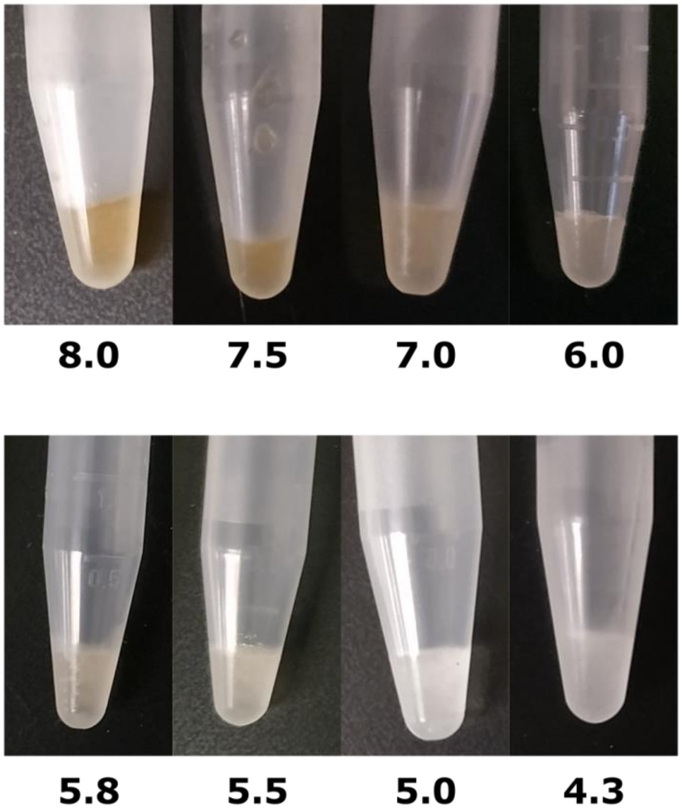


Supplementary Fig. 1. Humic acid minimization at different pH levels.

A decrease in pH results in less humic acid being extracted, but nucleic acid is absorbed into the soil at the same time. The pH of 5.8 was found to have the lowest humic content and a relatively low rate of nucleic acid absorption by the soil.

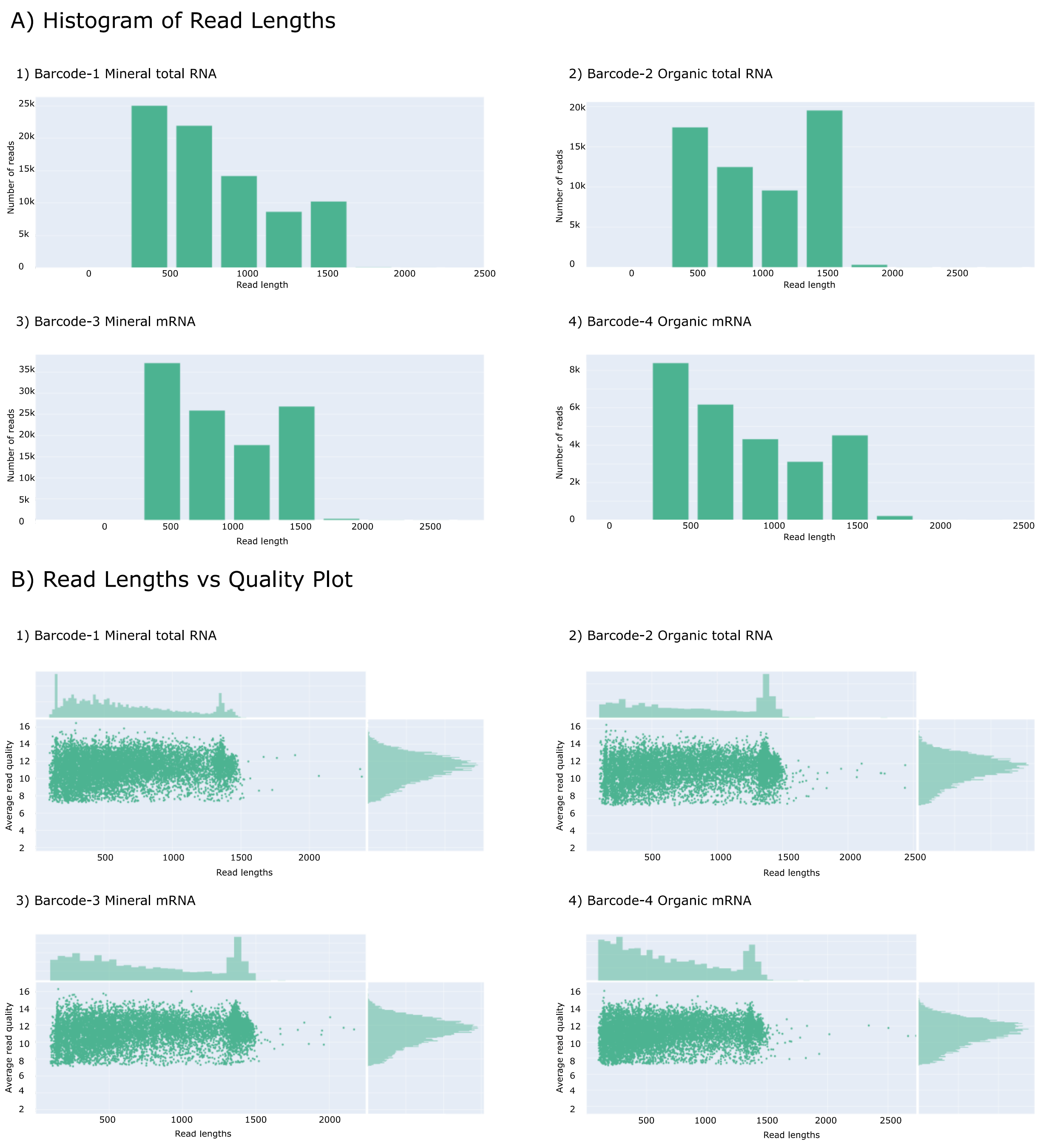


Supplementary Fig. 2. Histograms of read length and quality distributions of direct RNA sequencing data for mineral total RNA and organic total RNA as well as mineral mRNA and organic mRNA. Variations in single read length range from 654 to 818 bp. NanoPlot software (De Coster et al. 2018) was used to visualize the data.

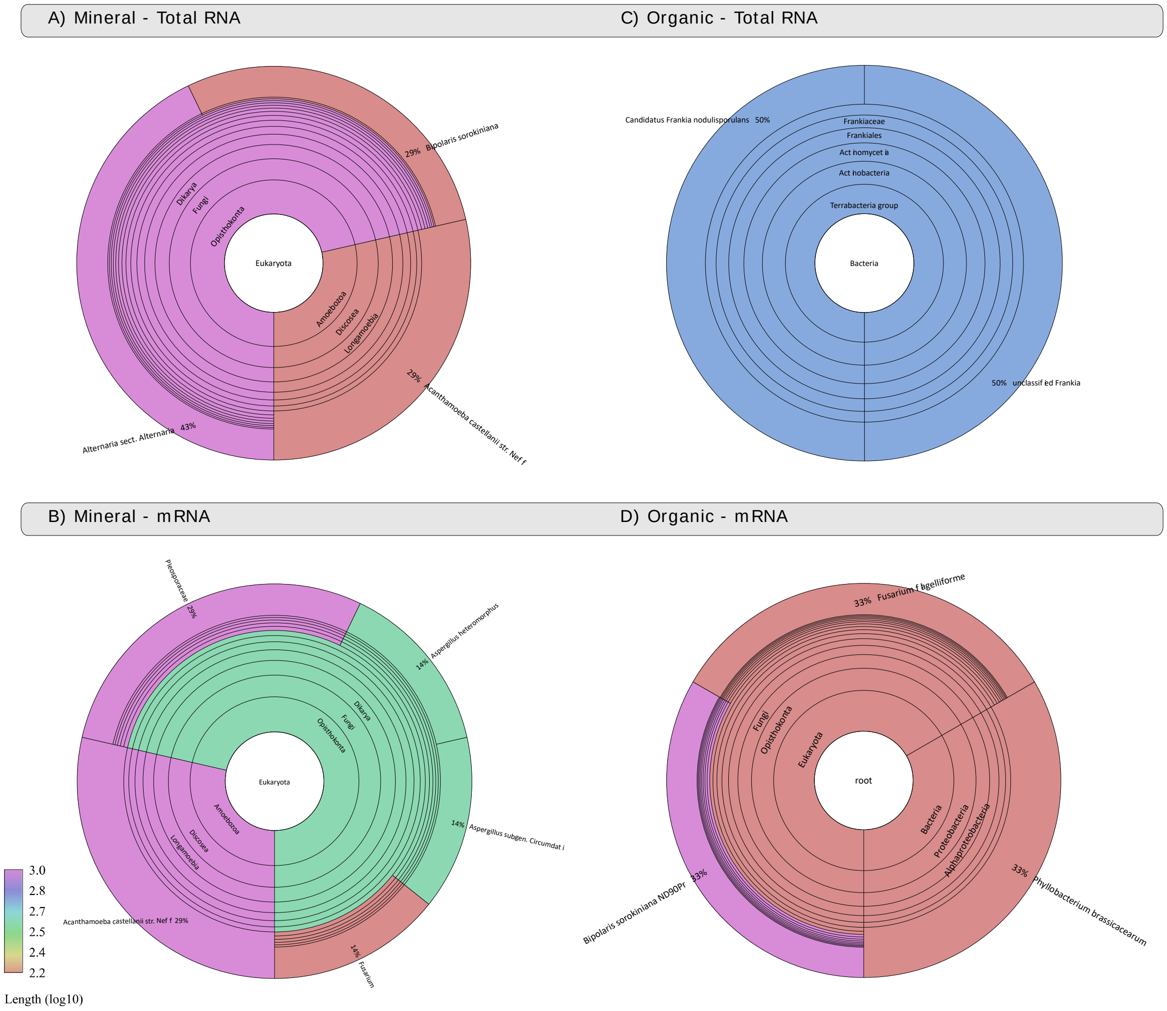


Supplementary Fig. 3. Taxonomical classification of unaligned clusters from Kraken2 step by Centrifuge. Mineral total RNA (A), mineral mRNAs (B), organic total RNA (C) and organic mRNA. The comparative analysis and visualization of results were done by Recentrifuge. Based on its developer's suggestion for nanopore sequencing reads, the LOGLENGTH was considered as a scoring scheme (Martí 2019).

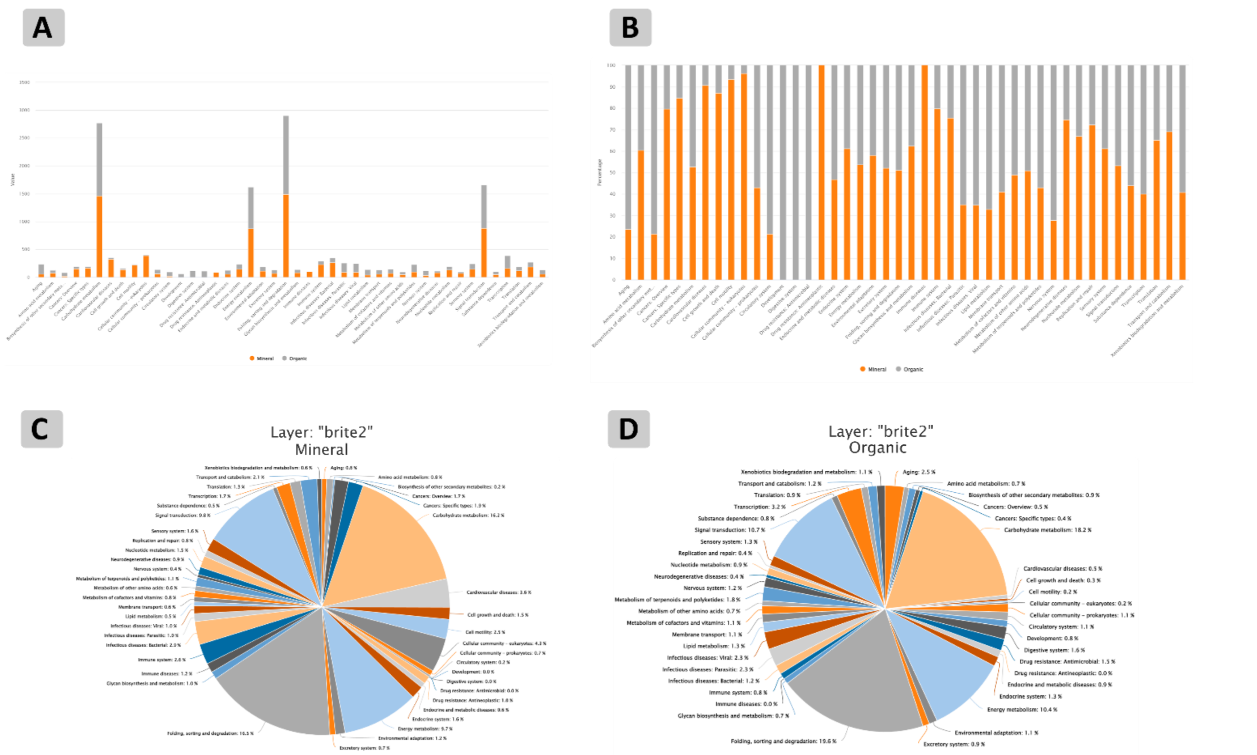


Supplementary Fig. 4. KEEG BRITE 2 analysis. Stacked barplots and pie charts of mineral and oganic soil clusters involved in KEGG BRITE 2. (A and B) Stacked barplots of mineral and organic data series before and after the normalization step. (C and D) Pie charts of mineral and organic clusters.

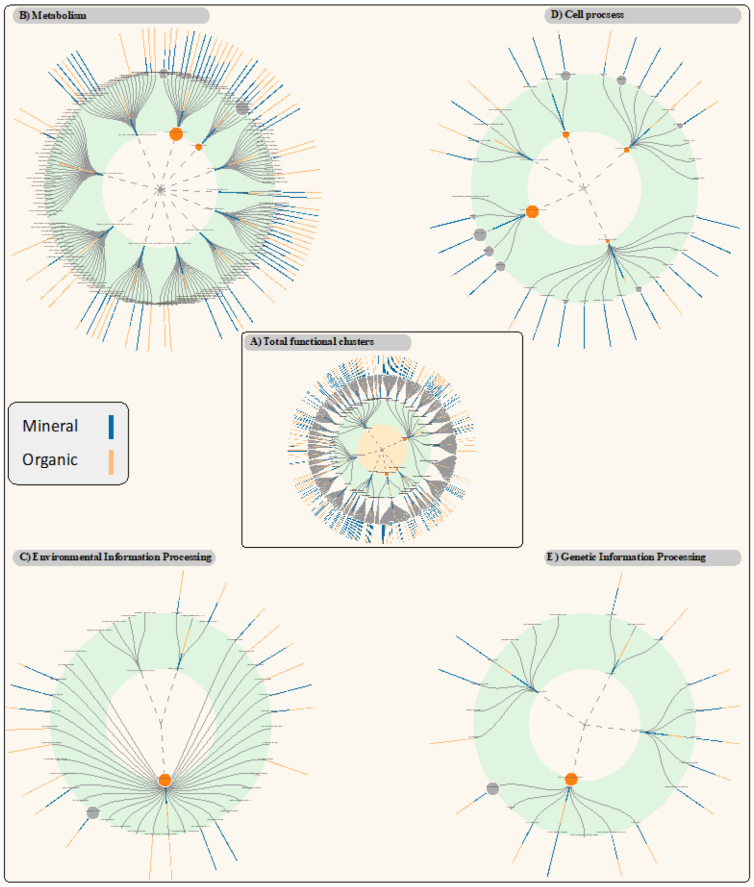


Supplementary Fig. 5. Circular dendogram of functional clusters in mineral and organic soil transcriptomes using FuncTree 2 which illustrated the hierarchical relationship of known biological functions in the KEGG database for mineral (blue) and organic (orange) metatranscriptomes. Each layer of the tree corresponds to each functional category: KEGG BRITE 1 (orang), KEGG BRITE 2 (gray), KEGG Pathway (blue).

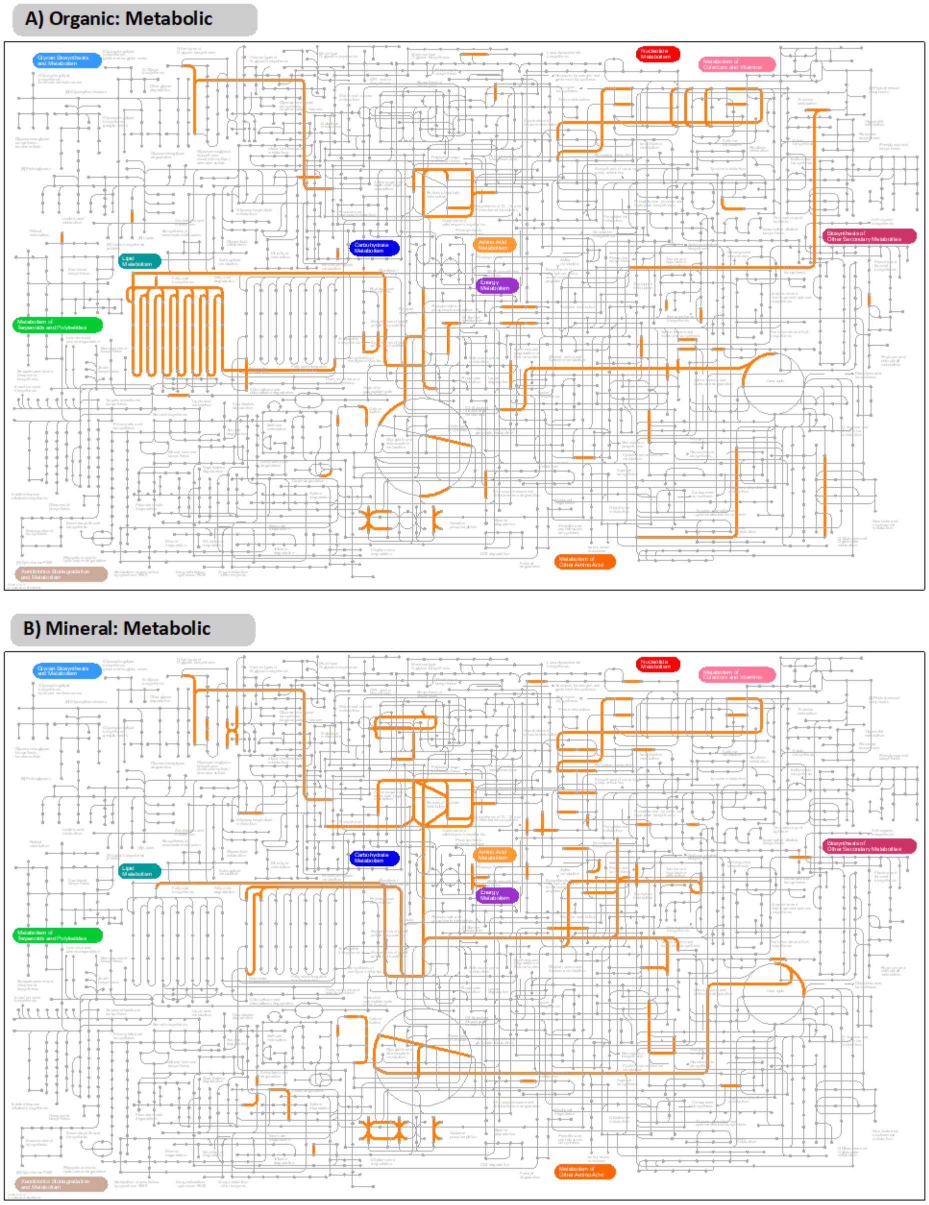


Supplementary Fig. 6. Metabolic and microbiome pathways raw input KOs of mineral and organic clusters mapped on metabolic and microbiome pathways by iPath3 (Darzi et al. 2018).

**Reference:**

Darzi, Y., Letunic, I., Bork, P., and Yamada, T. 2018. iPath3.0: interactive pathways explorer v3. Nucleic Acids Research 46:W510-W513.

De Coster, W., D’Hert, S., Schultz, D. T., Cruts, M., and Van Broeckhoven, C. 2018. NanoPack: visualizing and processing long-read sequencing data. Bioinformatics 34:2666-2669.

Martí, J. M. 2019. Recentrifuge: Robust comparative analysis and contamination removal for metagenomics. PLOS Computational Biology 15:e1006967.
