## Supplementary HTML Graphs for "Soil metatranscriptomics: An improved RNA extraction method toward functional analysis using nanopore direct RNA sequencing": S3-Mineral_rRNA.html

 Javascript must be enabled to view this page.

countunassignedtidrankscorek2\_alignedTR1\_PL 4311no\_rank2.83332superkingdom2.81511224phylum2.7441236class2.84128211class2.833356order2.86328216class2.63180840order2.722119060family2.9311783270no\_rank2.8268336no\_rank3.022976phylum3.01211783272no\_rank2.7311239phylum2.6291061class2.521385order2.522186817family2.582201174phylum2.8621760class2.82285009order3.1285011order2.722062family2.7221883genus2.792759superkingdom2.9733154no\_rank3.074751kingdom3.071451864subkingdom3.0614890phylum3.051716545no\_rank3.14147538subphylum3.04716546no\_rank3.04715989no\_rank3.04147550class3.04222543subclass3.041028384order3.04681950family3.045455genus3.042707350no\_rank3.0480884species3.044759273strain3.022698737no\_rank2.8233630no\_rank2.8225794phylum2.8
