## Supplementary HTML Graphs for "Soil metatranscriptomics: An improved RNA extraction method toward functional analysis using nanopore direct RNA sequencing": S4-Organic_rRNA.html

 Javascript must be enabled to view this page.

countunassignedtidrankscorek2\_alignedTR2\_PL 701no\_rank2.85742superkingdom2.81611224phylum2.7641236class2.621692040order2.5221692041family2.55128211class2.944356order2.9228216class3.02280840order3.0268525subphylum2.42228221class2.42257723phylum2.8811783257no\_rank2.87203682phylum2.87203683class2.82112order2.722126family2.7312691354order2.9222691359family2.822691355order2.6221914233family2.631783270no\_rank2.7368336no\_rank2.733976phylum2.72421783272no\_rank2.9811239phylum2.83191061class2.8221385order2.72186801class2.622186802order2.622909932class2.933200795phylum3.0114201174phylum2.9771760class2.91332759superkingdom2.88133154no\_rank2.874751kingdom2.87451864subkingdom2.874890phylum2.871716545no\_rank2.86147538subphylum2.86716546no\_rank2.861715989no\_rank2.85147550class2.831222543subclass2.821028384order2.82681950family2.825455genus2.822707350no\_rank2.8280884species2.822759273strain2.82222544subclass2.7225139order2.722698737no\_rank2.8233630no\_rank2.8225794phylum2.8
